# CD8+ T cell responses in convalescent COVID-19 individuals target epitopes from the entire SARS-CoV-2 proteome and show kinetics of early differentiation

**DOI:** 10.1101/2020.10.08.330688

**Authors:** Hassen Kared, Andrew D Redd, Evan M Bloch, Tania S. Bonny, Hermi Sumatoh, Faris Kairi, Daniel Carbajo, Brian Abel, Evan W Newell, Maria P. Bettinotti, Sarah E. Benner, Eshan U. Patel, Kirsten Littlefield, Oliver Laeyendecker, Shmuel Shoham, David Sullivan, Arturo Casadevall, Andrew Pekosz, Alessandra Nardin, Michael Fehlings, Aaron AR Tobian, Thomas C Quinn

## Abstract

Characterization of the T cell response in individuals who recover from SARS-CoV-2 infection is critical to understanding its contribution to protective immunity. A multiplexed peptide-MHC tetramer approach was used to screen 408 SARS-CoV-2 candidate epitopes for CD8+ T cell recognition in a cross-sectional sample of 30 COVID-19 convalescent individuals. T cells were evaluated using a 28-marker phenotypic panel, and findings were modelled against time from diagnosis, humoral and inflammatory responses. 132 distinct SARS-CoV-2-specific CD8+ T cell epitope responses across six different HLAs were detected, corresponding to 52 unique reactivities. T cell responses were directed against several structural and non-structural virus proteins. Modelling demonstrated a coordinated and dynamic immune response characterized by a decrease in inflammation, increase in neutralizing antibody titer, and differentiation of a specific CD8+ T cell response. Overall, T cells exhibited distinct differentiation into stem-cell and transitional memory states, subsets, which may be key to developing durable protection.

## Introduction

The emergence of severe acute respiratory syndrome coronavirus 2 (SARS-CoV-2) has rapidly evolved into a global pandemic. To date, over 35 million cases spanning 188 countries or territories have been reported with more than one million deaths attributed to coronavirus disease (COVID-19). The clinical spectrum of SARS-CoV-2 infection is highly variable, spanning from asymptomatic or subclinical infection, to severe or fatal disease ^1,2^. Characterization of the immune response to SARS-CoV-2 is urgently needed in order to better inform more effective treatment strategies, including antivirals and rationally designed vaccines.

Antibody responses to SARS-CoV-2 have been shown to be heterogenous, whereby male sex, advanced age and hospitalization status are associated with higher titers of antibodies ^3^. Low or even undetectable neutralizing antibodies in some individuals with rapid decline in circulating antibodies to SARS-CoV-2 after resolution of symptoms underscores the need to assess the role of the cellular immune response ^4^. Multiple studies suggest that T cells are important in the immune response against SARS-CoV-2, and may mediate long-term protection against the virus ^5–9^.

To date, studies that have evaluated SARS-CoV-2-specific T cells in convalescent individuals have focused on either characterization of responses to selected, well-defined SARS-CoV-2 epitopes, or broad assessment of T cell reactivity against overlapping peptide libraries ^6–10^. The assessment of the complete SARS-CoV-2 reactive T cell pool in the circulation remains challenging, and there is still much to be learned from capturing both the breadth (number of epitopes recognized) and depth of T cell response (comprehensive phenotype) to natural SARS-CoV-2 infection. A study by Peng *et al*. indicated that the majority of those who recover from COVID-19 exhibit robust and broad SARS-CoV-2 specific T cell responses ^8^. Further, those who manifest mild symptoms displayed a greater proportion of polyfunctional CD8+ T cell responses compared with severely diseased cases, suggesting a role of CD8+ T cells in ameliorating disease severity.

Many current COVID-19 vaccine candidates primarily incorporate the SARS-CoV-2 spike protein to elicit humoral immunity ^11–13^. However, whether these approaches will induce protection against SARS-CoV-2 infection, or COVID-19 remain unknown. Gaining insight into the immune response that is induced by natural SARS-CoV-2 infection will be key to advancing vaccine design. Specifically, there is a need to identify what T cells are targeting in the viral proteome, their functional characteristics, and how these might correlate with disease outcomes. In this study, our analytical strategy progressed beyond these earlier findings by identifying dozens of epitopes recognized by CD8+ T cells that spanned different viral proteins in COVID-19 convalescent subjects, and simultaneously revealed the unmanipulated phenotypic profiles of these cells. These new findings can be exploited to further guide epitope selection for rationally designed vaccine candidates and vaccine assessment strategies.

## Results

### SARS-CoV-2-specific CD8+ T cell response in COVID-19 convalescent donors is broad and targets the whole virus proteome

To study the SARS-CoV-2 specific CD8+ T cell repertoire in COVID-19 convalescent donors, a mass cytometry-based multiplexed tetramer staining approach was employed to identify and characterize (i.e. phenotype) SARS-CoV-2-specific T cells *ex vivo* (Figure 1A). A total of 30 convalescent plasma donors (confirmed by PCR at time of infection) with HLA-A*01:01, HLA-A*02:01, HLA-A03:01, HLA-A*11:01, HLA-A*24:02 and HLA-B*07:02 alleles were evaluated ^3^. The individuals included 18 males and 12 females ranging between 19 and 77 years old, and were a median of 42.5 days (interquartile range 37.5-48.0) from initial diagnosis (Table S1). The population was grouped into tertiles according to their overall anti-SARS-CoV-2 IgG titers, based on semi-quantitative ELISA results against SARS-CoV-2 S protein (Table S2). Additional plasma-derived parameters such as neutralizing antibody titers, inflammatory cytokines and chemokines were used to associate the cellular SARS-CoV-2-specific T cell response with the humoral and inflammatory response (Figure 1A). There was a strong correlation between the donors’ anti-S IgG levels and the neutralizing antibody activity (Fig S1A). Levels of some inflammatory mediators were associated with age, sex, neutralizing antibody activity and neutralizing antibody titers (Fig S1B-D).

**Figure 1.**
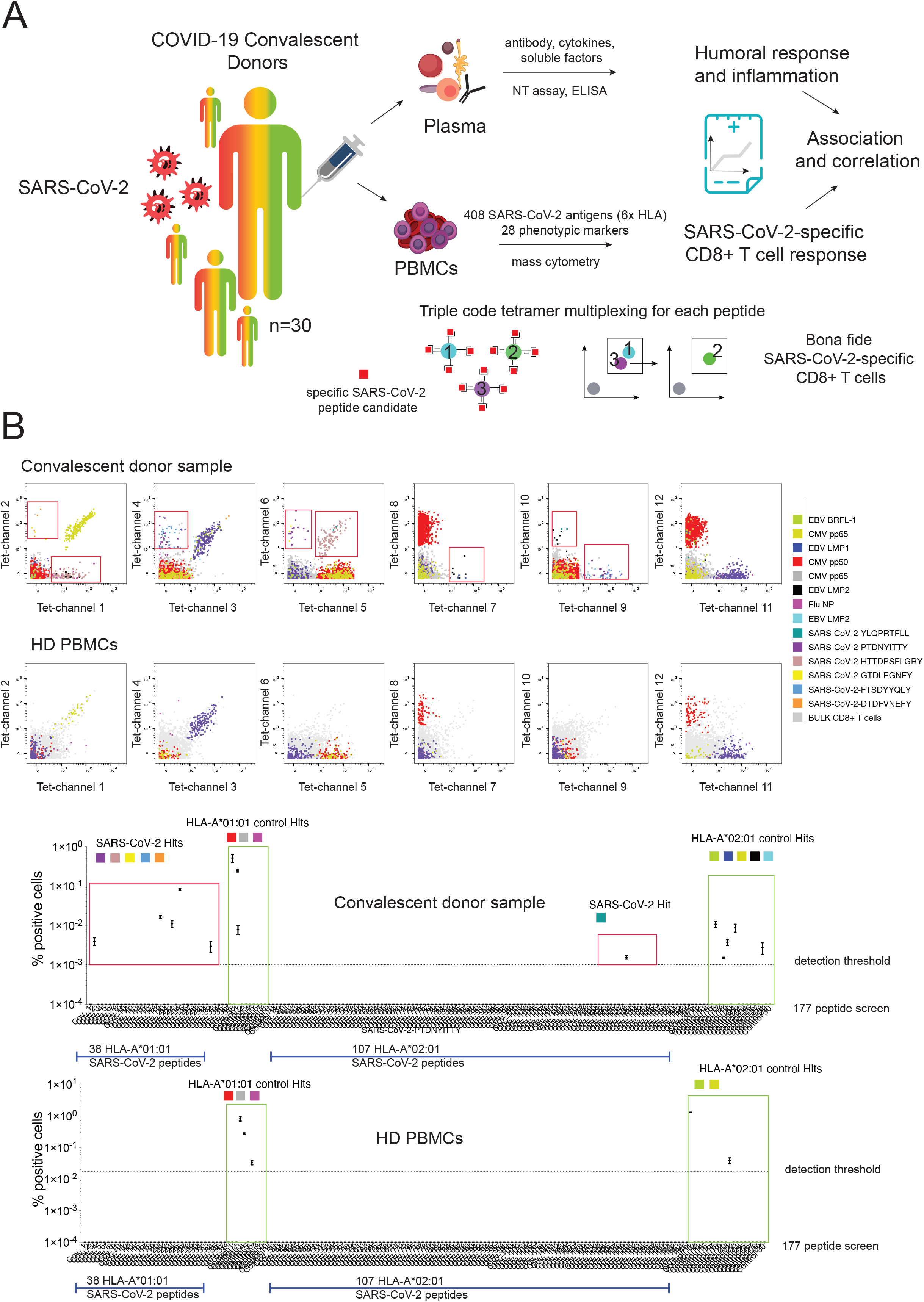
Identification and characterization of SARS-CoV-2-specific CD8+ T cells from SARS-CoV-2 convalescent donors. **A)** Visualization and schematic overview of the experimental workflow. SARS-CoV-2-specific CD8+ T cells were identified and simultaneously characterized in PBMCs from convalescent donors by screening a total of 408 SARS-CoV-2 candidate epitopes across six HLAs using a mass cytometry based highly multiplexed tetramer staining approach. Frequencies and phenotypic profiles of SARS-CoV-2-specific T cells were associated and correlated with the cross-sectional sample-specific humoral response and inflammation parameters. **B)** Representative staining and screening example for SARS-CoV-2-specific CD8+ T cells from a convalescent donor sample. Shown is a screen probing for 145 SARS-CoV-2 candidate antigens (HLA-A02 and HLA01) and 31 SARS-CoV-2 unrelated control antigens. Healthy donor PBMCs were run in parallel. Red boxes indicate SARS-CoV-2-specific T cell hits. Screening data shows the values and means from the 2 technical replicates (2 staining configurations). Bona fide antigen-specific T cells were defined based on different objective criteria set (Methods).

Hundreds of candidate epitopes spanning the complete SARS-CoV-2 genome were recently identified as potential targets for a CD8+ T cell response to SARS-CoV-2 ^14,15^. A triple-coded multiplexed peptide-MHC tetramer staining approach was used to screen 408 potential epitopes for recognition by T cell responses across 6 different HLA alleles: HLA-A*01:01, HLA-A*02:01, HLA-A03:01, HLA-A*11:01, HLA-A*24:02 and HLA-B*07:02 ^16,17^. In addition, CD8+ T cells were probed for reactivity against up to 20 different SARS-CoV-2-unrelated control peptides per HLA for each sample (CMV-, EBV-, Influenza-, Adenovirus-, and MART-1-derived epitopes; Table S3). The detection of bona fide antigen-specific T cells was based on the assessment of several objective criteria such as signal versus noise, consistency between two technical replicates, and detection threshold. In this study, an average limit of detection of 0.0024% (bootstrapping confidence interval of 0.0017 and 0.005 under a confidence level of 95%) was achieved for antigen-specific T cells. Depending on the individual’s HLA allele repertoire, between 48 and 220 peptides were simultaneously screened per participant.

Figure 1B shows an example of the identification of antigen-specific CD8+ T cells in a COVID-19 convalescent donor screened for a total of 145 SARS-CoV-2 antigen candidates and 32 common (SARS-CoV-2 unrelated) control antigens across two HLA alleles. CD8+ T cells reactive to six different SARS-CoV-2 epitopes and eight control antigens were detected, including peptides derived from Influenza (FLU), Epstein Barr Virus (EBV), and Cytomegalovirus (CMV). In parallel, commercially obtained healthy donor PBMCs were run and similar common virus antigen specificities were identified. Notably, SARS-CoV-2 specific CD8+ T cells were not detected in any of the healthy donors recruited before the official SARS-CoV-2 pandemic (n=4).

Amongst all 408 SARS-CoV-2 peptide candidates tested, 52 unique peptide reactivities (hits) were detected which were distributed across a total of 132 SARS-CoV-2 peptide epitopes (Fig. 2A). Almost all individuals screened demonstrated a CD8+ T cell response against SARS-CoV-2 (29/30), and individual hits ranged from 0 to 13 with >40% of all individuals showing more than five different SARS-CoV-2 specificities. The frequency of these cells ranged from 0.001% to 0.471% of total CD8+ T cells (Table S4). In addition, a total of 130 T cell hits against common control peptides were detected in these donor samples (0.001%.to 1.074% of total CD8+ T cells) (Table S4). Interestingly, the majority of unique T cell hits were directed against epitopes associated with non-structural proteins such as nsp, PLP and ORF3a (Fig 2B). Of all the hits that were detected in the cross-sectional sample, the most common reactivities were against spike (structural, 23.02%) and ORF3a (non-structural, 19.42%). By contrast, nucleocapsid-specific CD8+ T cells had significantly higher frequencies as compared to spike- or non-structural protein-specific T cells (Fig. 2C). The total number of recognized epitopes was distributed differently across the individual HLA alleles that were tested (Fig 2D and Fig S2), whereby T cell responses were identified against six to 14 different epitopes per allele (Fig 2E). For the purpose of the study, events detected in at least three donor samples or in more than 35% of donors for each allele group were defined as SARS-CoV-2 high-prevalence epitope hit responses.

**Figure 2.**
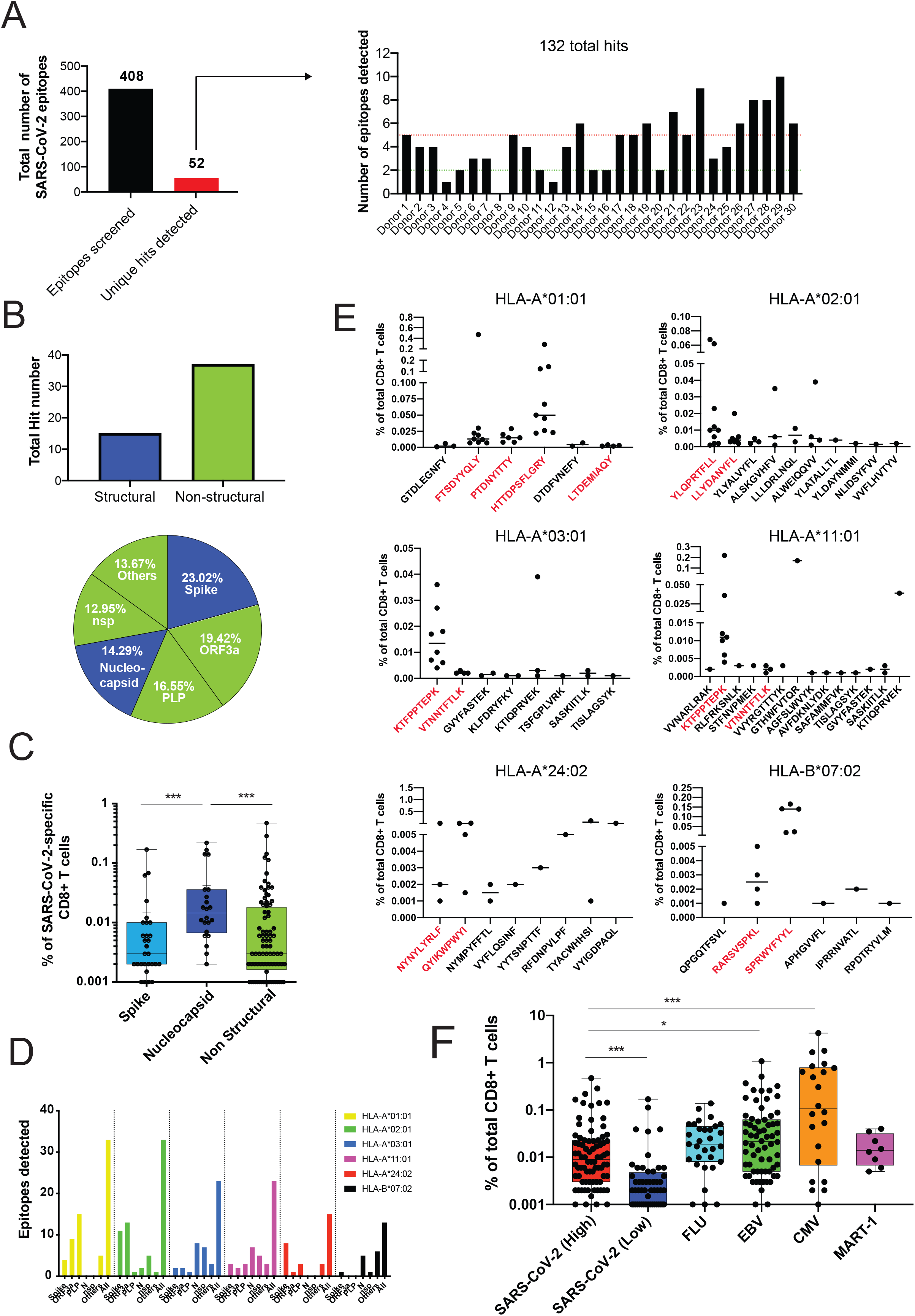
Breadth and magnitude of SARS-CoV-2-specific CD8+ T cells. **A)** Bar plots summarizing the absolute numbers of SARS-CoV-2 antigen specificities detected across donors within cross-sectional sample. Out of 408 SARS-CoV-2 peptide candidates 52 unique peptide hits were detected. Between 0 and 13 unique hits were detected in each donor sample (five or more hits in >40% of all donors). In total, 132 SARS-CoV-2-specific T cell hits were detected **B)** Delineation of T cell reactivities against the SARS-CoV-2 proteome. The majority of epitope hits detected derived from non-structural SARS-CoV-2 proteins. Pie chart displaying the percentages of epitopes detected derived from structural (Nucleocapsid, Spike) and non-structural (nsp, PLP, ORF3a, others) proteins spanning the full proteome of SARS-CoV-2. **C)** Frequencies of SARS-CoV-2 specific T cells reactive with epitopes derived from spike, nucleocapsid and non-structural proteins. Highest frequencies were detected for T cells targeting peptides from the nuleocapsid protein. Each dot represents one hit. **D)** Numbers of epitopes from the different protein categories detected across all six HLA alleles tested. **E)** Definition of high- and low-prevalence hits per HLA allele. Plots showing individual peptide hits for each allele. Each dot represents one hit. High-prevalence epitope hits are indicated in red and were defined as events detected in at least three donor samples or in more than 35% of donors for each allele group. **F)** Comparison of frequencies of SARS-CoV-2 specific T cell and T cells reactive with influenza, EBV, CMV, or endogenous MART-1 epitopes. The percentage of SARS-CoV-2 specific T cells was higher for epitopes categorized as high prevalence hits but lower than the frequencies of T cells reactive with EBV or CMV antigens detected. *p<0.1, **p<0.01, *** p<0.001. Kruskal-Wallis test. p-values were adjusted for multiple testing using the Benjamini-Hochberg method to control the false discovery rate.

Based on these criteria, at least two peptides per HLA allele were defined as high-prevalence response hits (Fig 2E). Of note, the frequencies of high-prevalence SARS-CoV-2-specific T cells were significantly higher as compared to their low-prevalence counterparts (Fig 2F). Frequencies of high-prevalence SARS-CoV-2-specific T cells were similar to those of FLU-specific T cells detected in the same cross-sectional sample, but significantly lower than frequencies of T cells reactive for EBV or CMV peptides (Fig 2F). In summary, these data show a reliable detection of multiple SARS-CoV-2 T cell hits and indicate a broad recognition of epitopes by CD8+ T cell responses against the SARS-CoV-2 proteome during recovery from COVID-19.

### SARS-CoV-2-specific CD8+ T cells exhibit a unique phenotype and can be classified into different memory subsets

Our multiplexed tetramer staining approach enables deep phenotypic characterization of antigen-specific T cells. By using a panel comprising 28 markers that were dedicated to T cell identification and profiling, including several markers indicative of T cell differentiation (Table S5), the phenotypic profiles of all SARS-CoV-2-specific T cells detected in this cross-sectional sample were further analysed.

To compare the phenotypes of antigen-specific T cells targeting different SARS-CoV-2 proteins, the frequencies of T cells expressing all markers were determined (Fig 3A). Despite some phenotypic heterogeneity, the majority of SARS-CoV-2-specific T cells grouped together and were distinct from T cells that were specific for CMV-, EBV-, or FLU-derived epitopes detected in the same samples; the same outcome was reached when displaying the data as a two-dimensional UMAP plot (Fig 3B). SARS-CoV-2 specific T cells showed an intermediate phenotype between MART-1-specific T cells, which are predominantly naïve (CCR7 high and CD45RA high), and memory FLU-specific T cells ^18^.

**Figure 3.**
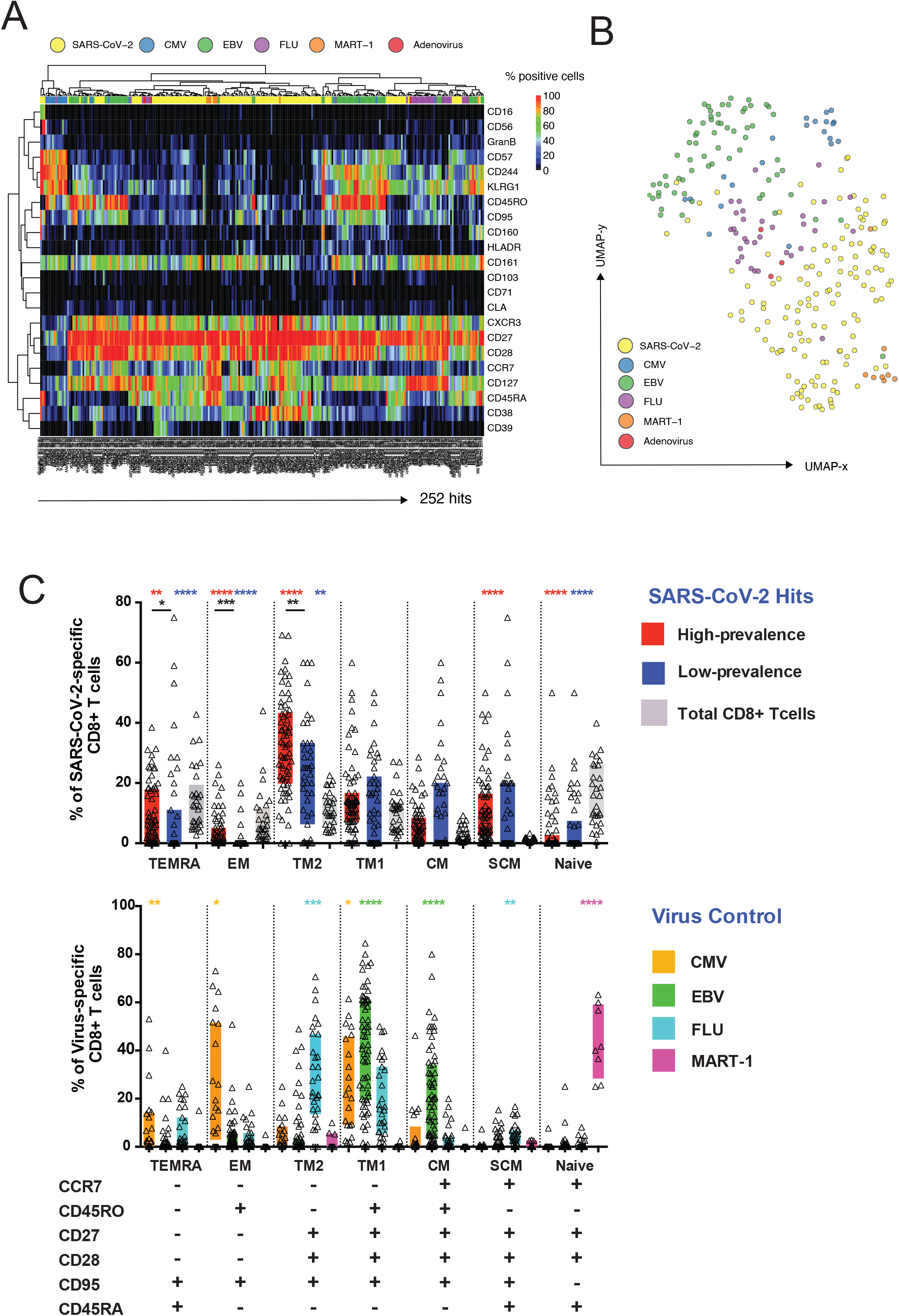
SARS-CoV-2-specific CD8+ T cells display a unique phenotype and can be categorized into different subsets. **A)** Heatmap summarizing the expression frequencies of all phenotypic markers analyzed among the total pool of SARS-CoV-2-specific and unrelated control antigen-specific CD8+ T cells detected in the same cross-sectional sample. The majority of SARS-CoV-2 specific T cells clusters differently from common virus-specific T cells. Antigen-specificities and phenotypic markers were clustered using Pearson correlation coefficients as distance measure. **B)** UMAP plot showing the clustering of all antigen-specific T cells by antigen category. SARS-CoV-2-specific CD8+ T cell occupy the lower region of the two-dimensional map. Clustering is based on the expression of all phenotypic markers assessed. Each dot represents one hit. **C)** Differentiation profiles of SARS-CoV-2-specific CD8+ T cells and common virus control antigen-specific T cells. Based on the expression of the markers below the bar diagrams, antigen-specific and total CD8+ T cells were categorized into distinct states of differentiation. SARS-CoV-2-specific T cells were enriched in TSCM and TM2 cells. Control virus hits could be separated into distinct subsets dependent on the target epitope. *p<0.1, **p<0.01, *** p<0.001. Wilcoxon rank sum test. TSCM (stem-cell memory cells), TM (transitional memory cells), TEMRA (terminal effector memory cells re-expressing CD45RA), EM (effector memory cells), CM (central memory cells).

An early differentiated memory phenotype has recently been described for SARS-CoV-2-specific T cells ^9^. SARS-CoV-2-specific T cells were separated into subpopulations based on the stages of T cell differentiation, further split into high- and low-prevalence response hits as earlier defined, and their frequencies compared with one another, as well as with total CD8+ T cells. Likewise, these were compared with the differentiation profiles of T cells reactive against common virus antigens and MART-1. The classification into functionally different T cell subsets following antigen encounter is based on the expression of different marker combinations, which describe a progressive T cell differentiation and allow to delineate a dynamic transition between memory and effector cell function ^19^ (Fig 3C and S3). When compared to the total CD8+ T cell population, SARS-CoV-2-specific T cells were significantly enriched for cells with stem-cell memory (SCM) and transitional memory cells 2 (TM2) phenotypes. More specifically, high-prevalence SARS-CoV-2-specific T cells were skewed toward a phenotype that is typical of terminal effector memory cells re-expressing CD45RA (TEMRA), effector memory cells (EM) and TM2 cells, while their low-prevalence counterparts were enriched with SCM and central memory (CM) cells. In contrast, MART-1-specific T cells were naïve, FLU-specific T cells were predominantly of a TM2 phenotype, EBV-specific T cells were largely characterized by TM1 and CM phenotypes, and CMV-specific T cells were more differentiated as reflected by a strong effector component.

### Expansion of highly differentiated SARS-CoV-2-specific CD8+ T cells in convalescent donors

To gain further insight into the phenotypes of SARS-CoV-2-specific CD8+ T cells, the expression of all the phenotypic markers were compared between T cells exhibiting high-with those exhibiting low-prevalence epitope responses. Similar to our findings in the total pool of SARS-CoV-2-specific CD8+ T cells, a heterogenous marker expression was detected across these cells, but no specific clustering with respect to the epitope response prevalence (Fig S4A). To further compare the phenotypes of T cells from high-vs. low-prevalence epitope response categories, the high-dimensionality of the dataset was reduced and the phenotypic information plotted from Figure S4A using principal component analysis (PCA) (Fig S4B). The PCA displayed a skewing of high-prevalence SARS-CoV-2-specific T cells towards late T cell differentiation (CD57 and CD161), in contrast to the low-prevalence response hits characterized by early differentiation markers (CD27, CD28, CCR7). In order to quantify this spatial distribution, the individual expression of all markers was evaluated and the frequencies for each marker compared between the high- and low-prevalence response hits. Significantly higher frequencies of T cells expressing CD57 and Granzyme B were detected amongst high-prevalence SARS-CoV-2-specific T cells, while the frequencies of CCR7-expressing cells were substantially higher amongst the low-prevalence hit responses (Fig 4A). These findings were further confirmed when overlaying the SARS-CoV-2-specific T cells on a two-dimensional UMAP plot created based on the full phenotypic panel (Fig 4B). The majority of T cells that had been categorized as high-prevalence response hits were associated with the expression of CD57 and Granzyme B, while their low-frequency counterparts detected in the same donors were characterized by a high CCR7 expression.

**Figure 4.**
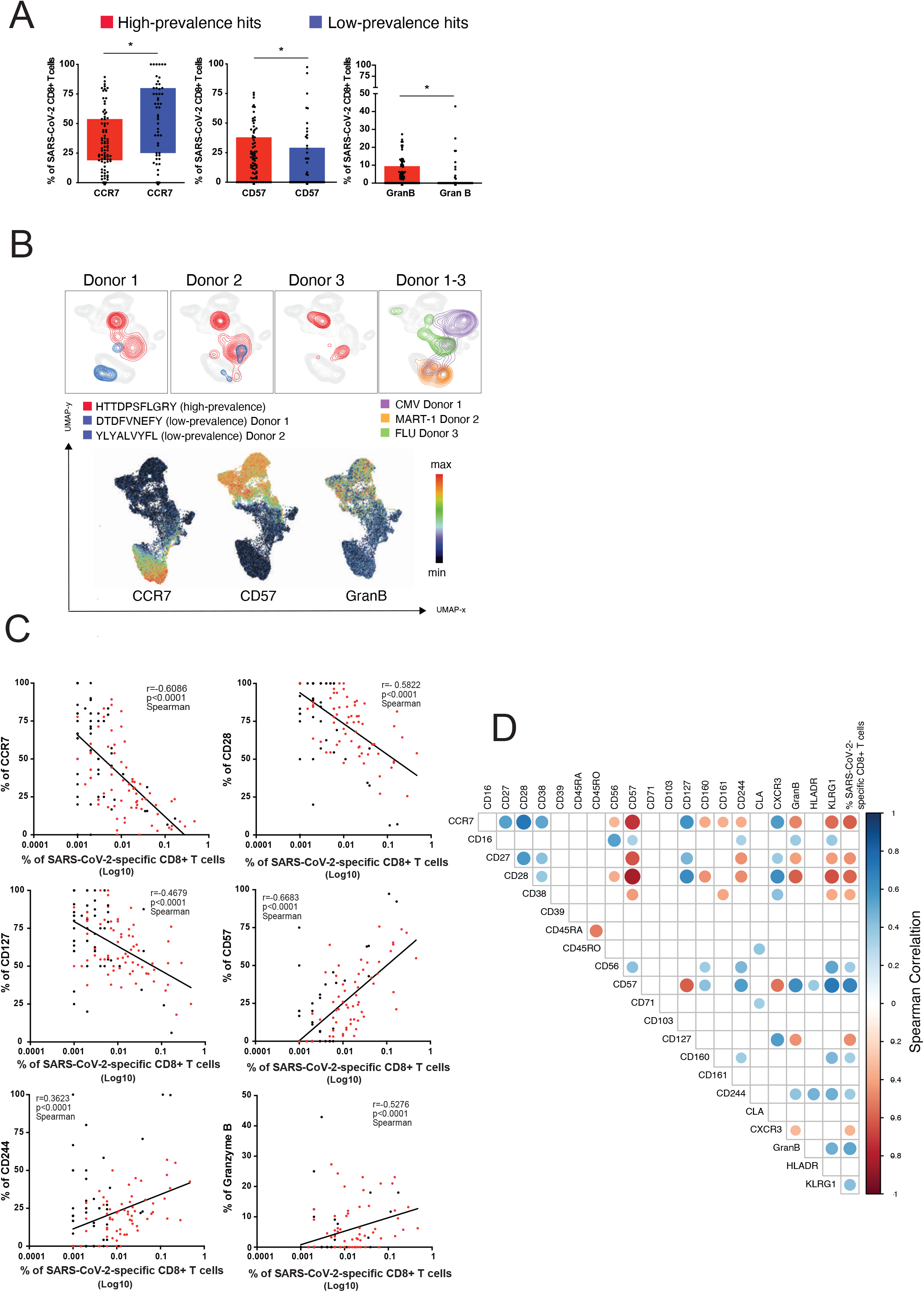
Expansion of highly differentiated SARS-CoV2-specific CD8+ T cells in convalescent donors. **A)** Boxplots showing differences in the expression of markers between high- and low-prevalence response hits. High-prevalent response hits showed a higher expression of markers associated with differentiation. Each dot represents one donor. *p<0.1, **p<0.01, *** p<0.001. Kruskal-Wallis test. p-values were adjusted for multiple testing using the Benjamini-Hochberg method to control the false discovery rate. UMAP plot showing the relative position of high- and low-prevalence response hits in the high-dimensional space. Data from 3 donors is shown. **C)** Scatterplots showing the correlations between SARS-CoV-2-specific T cell frequencies and differentiation marker expression. The magnitude of antigen-specific T cells correlated with the expression of markers associated with T cell differentiation. The correlations were calculated with the Spearman’s rank-order test. Red dots are high prevalence response hits. **D)** Correlogramm showing the correlation between all phenotypic markers and frequencies of SARS-CoV-2-specific T cells. Later stage differentiation markers positively correlated with higher frequency SARS-CoV-2-specific T cells. Spearman’s correlation coefficients were indicated by a heat scale whereby blue color shows positive linear correlation, and red color shows negative linear correlation. Only significant correlations are shown (*p<0.05, p-values were adjusted for multiple testing using the Bonferroni method).

High-prevalence SARS-CoV-2-specific T cells were detected at a higher frequency (Fig 2F) as compared to their low-prevalence counterparts. Therefore, assessment of the magnitude of the SARS-CoV-2-specific T cell response was also correlated with their phenotypes. Interestingly, a negative correlation between the frequency of SARS-CoV-2 specific CD8+ T cells and the expression of markers associated with early T cell differentiation was observed (CD28, CCR7, CD127, CD27, CD38, and CXCR3) (Fig. 4C and 4D). In contrast, the level of expression of markers that are associated with late stage T cell differentiation (CD244, CD57, Granzyme B, and KLRG1) correlated positively with increasing frequencies of SARS-CoV-2 specific CD8+ T cells.

### Time-dependent evolution of SARS-CoV-2-specific CD8+ T cell response, inflammation and humoral immune response

To examine the relationship between inflammation, humoral immunity, and the T cell response, the frequencies of SARS-CoV-2-specific CD8+ T cells were evaluated against their IgG to Spike titer and neutralizing antibody activity (measured by NT AUC) (Fig 5A). Interestingly, although the phenotypic clustering of SARS-CoV-2-specific CD8+ T cells was not associated with IgG titer tertiles (Fig S4A), NT AUC correlated negatively with expression of markers associated with an immature or early differentiated phenotype (CCR7, CD28, CD45RA, CD127, CXCR3), while correlating positively with CD57 and CD161 (Fig 5A-B). Next, assessment of the association between inflammatory molecules and SARS-CoV-2-specific T cells was conducted. Inflammation can indirectly regulate the persistence of antigen-specific T cells in the absence of TCR stimulation or during chronic infection by modulating the homeostatic cytokine profile ^20,21^. Overall, the correlation between inflammatory mediators and the expression of individual markers on SARS-CoV-2-specific T cells, or the T cell frequency, remained weak (Fig 5A). Finally, the evolution of the SARS-CoV-2-directed T cell response against time based on the last detection of SARS-CoV-2 specific mRNA was modeled in each donor (Table S1). Interestingly, an increase in the breadth of the specific CD8+ T cell response was observed during the resolution phase of the disease, peaking at approximately six weeks (Fig S5). Longer recovery time was associated with higher frequencies of cells expressing markers of terminal T cell differentiation (CD57, CD244 and KLRG-1) and activation (HLA-DR), indicating a positive correlation between recovery time and T cell maturation (Fig 5A and 5C). Plasma levels of several cytokines (IL-18, TARC, MCP-1, VEGF) also decreased over time suggesting a negative correlation between recovery time and inflammation (Fig S1A).

**Figure 5.**
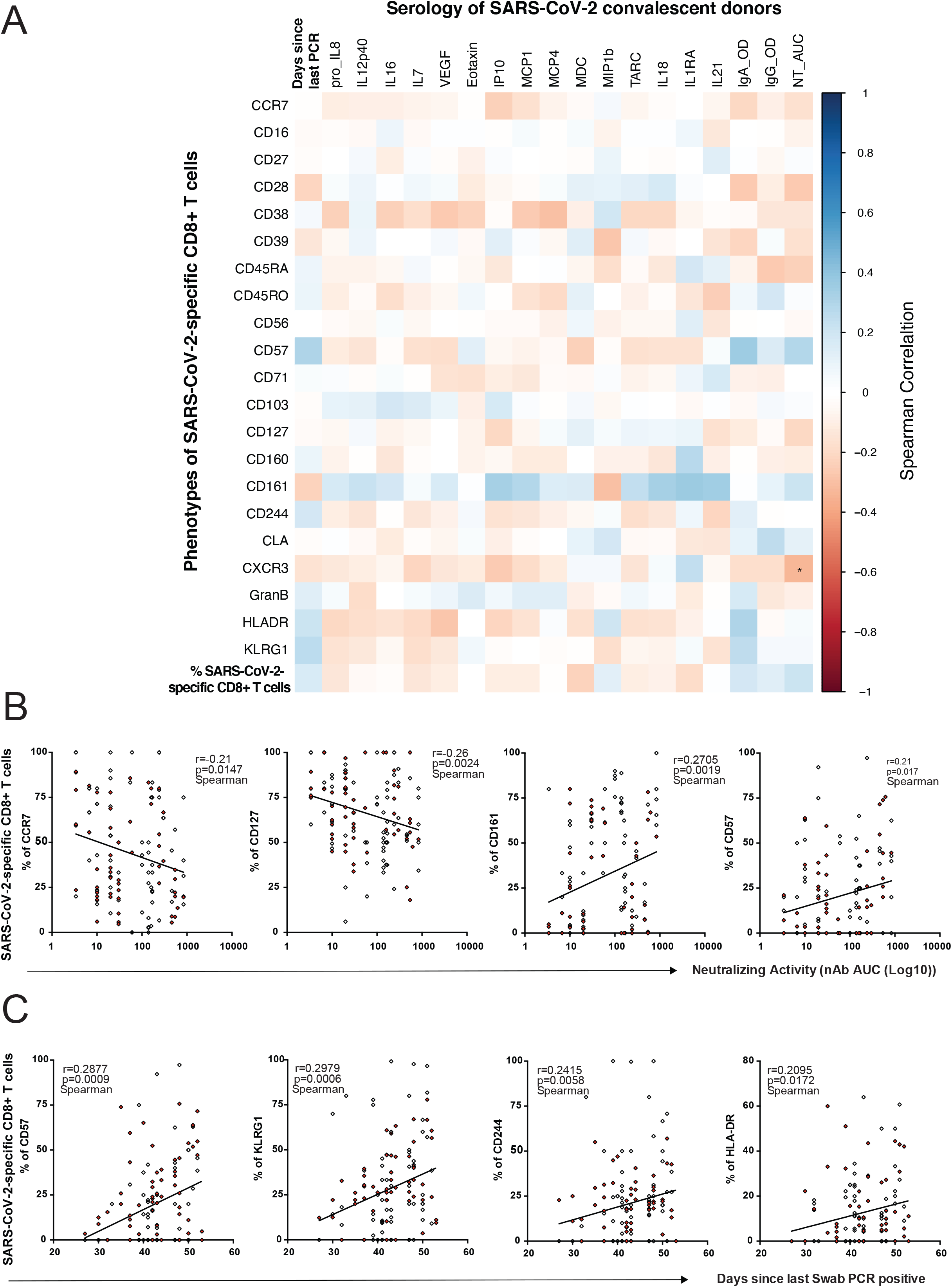
Time-dependent evolution of SARS-CoV-2-specific CD8+ T cell response, inflammation and humoral immune response. **A**) Correlation matrix showing the associations between frequencies and phenotypic markers of SARS-CoV-2-specific T cells and serological markers and recovery time (days since PCR). Spearman correlation (blue: positive correlation, red: negative correlation). *p<0.05, p-values were adjusted for multiple testing using the Bonferroni method. **B)** Scatterplots showing the correlations between marker expression on SARS-CoV-2-specific T cells and neutralizing antibody activity. Higher expression of markers associated with T cell differentiation was associated with a stronger neutralizing antibody activity. **c)** Scatterplots showing the correlations between marker expression on SARS-CoV-2-specific T cells and recovery time. The expression of markers associated with late stage differentiation correlated with the donors’ recovery time (days since last swab PCR positive). Correlations were calculated with the Spearman’s rank-order test. Red dots indicate high-prevalence response hits.

These data suggest that during early recovery from COVID-19, an overall, time-dependent decrease in inflammation is associated with sustained and effective antibody neutralizing activity with progressive differentiation of a broad and functional SARS-CoV-2-specific CD8+ T cell response (Fig S6).

## Discussion

An improved understanding of natural immunity to SARS-CoV-2 is needed to advance development of prevention strategies and/or treatment options for COVID-19. Recent findings suggest that T cells confer protection, whereby virus-specific memory T cell responses have been demonstrated in the majority of those who recover from COVID-19 even in the absence of detectable circulating antibodies ^9^. Moreover, the detection of T cells that are specific for the original SARS-CoV nucleoprotein in patients years after infection highlights the potential role of T cells in generating long lasting immunity against the virus ^7^. A mass cytometry based peptide-MHC-tetramer staining strategy ^17^ was applied, whereby 408 SARS-CoV-2 candidate epitopes were screened spanning 6 different HLA alleles. This enabled an *ex vivo* identification and true phenotypic characterization of virus-specific T cells in COVID-19 convalescent individuals without an *in-vitro* culture or stimulation bias which could affect the cellular phenotype, in contrast to prior studies using overlapping peptide pools ^6–8^. The high detection rate of SARS-CoV-2-specific CD8+ T cells across these COVID-19 convalescent donors is consistent with previous reports ^6–8^. In addition to the detection of epitopes previously described by others, over a third (i.e. 35%) of the antigen-specific T cells identified here have not been previously reported (Table S6), thereby highlighting the sensitivity of the adopted screening approach ^8,9,22–27^. However, given the low frequencies of many of these CD8 T+ cells, it is possible that these were below the detection threshold since T cell counts are very low in acutely infected patient samples ^28^. The T cell response in our study was directed against the full SARS-CoV-2 proteome with the majority of CD8+ T cells targeting epitopes derived from internal and/or non-structural virus proteins, which is in agreement with the recent findings by others ^8^. Moreover, half of the high-prevalence response hits identified for each HLA comprised antigens derived from non-structural proteins. In total, 12 highly-prevalent SARS-CoV-2 specific CD8+ T cell responses were identified, several of which overlapped with the immunodominant peptides detected by others ^8^, while some differed by the HLA type or the viral proteins that were assessed. The overall breadth and magnitude of the SARS-CoV-2-specific CD8+ T cell response may depend on the viral load, the severity of the disease and the intensity of the priming during the acute phase of the disease. Therefore, our collective findings support inclusion of a broad repertoire of SARS-CoV-2 epitopes in future vaccine designs ^8^.A unique phenotype for SARS-CoV-2-specific T cells was observed that was distinct from other common virus-specific T cells detected in the same cross-sectional sample. In particular, an enrichment in cells with a stem-cell and transitional memory phenotype was observed as compared to total and other virus-specific T cells. The inclusion of CD27, CD28 and CD95 in the evaluation of T cell differentiation states facilitates a granular characterization of T cell progression and reveals memory and effector capabilities of these cells. In addition, we observed a broad expression of CD127 across all hits detected, inferring proliferative functionality. A similar early differentiated memory phenotype has recently been described for SARS-CoV-2-specific T cells, and was further characterized by polyfunctionality and proliferative capacity ^9,29^. The potential of T_SCM_ to differentiate into various T cell memory subsets might contribute to durable protection against SARS-CoV-2 in COVID-19 convalescent donors. However, Kared *et al*., recently showed that an alteration in the Wnt signaling affects the differentiation capabilities of T_SCM_ in older patients ^30^.

In the current study, no significant impact of age was observed on the quantity or quality of SARS-CoV-2-specific T cells. The potential role of T_SCM_ in SARS-CoV-2 immune protection remains to be assessed in larger cohorts with longitudinal follow-up studies. Higher T cell frequencies were observed in high-prevalence epitope responses, and an increased expression of late differentiation markers (CD57, Granzyme B) vs. early differentiation markers (CCR7) was observed in high-versus low-prevalence epitope responses, respectively. Overall, the increased expression of markers associated with T cell differentiation correlated with the frequencies of SARS-CoV-2-specific T cells detected in this cross-sectional sample. The evolving profiles of epitope-specific responses during the resolution phase of the disease suggest a continuous proliferation and dynamic differentiation of T_SCM_ into effector memory CD8+ T cells. Previous studies described an activated phenotype across SARS-CoV-2-specific T cells during acute COVID-19 infection ^9^, which was not consistently observed in our cross-sectional sample. The quiescent phenotype observed in our study may be a consequence of down-regulated activation after viral clearance, homeostatic proliferation associated with the resolution of inflammation, or a combination of both. Indeed, while there was a decrease in inflammation in this cross-sectional sample over time, the expansion and progressive differentiation of a broad and functional SARS-CoV-2-specific T cell response correlated with the neutralizing antibody activity and donor recovery time. Our findings bring new insights into the viral targets and dynamics of the SARS-COV-2-specific CD8+ T cell response. Nevertheless, it remains to be investigated whether a T cell response to a broad diversity of epitopes is relevant at the early and acute stages of the disease, and whether they have a protective role at the primary site of infection, as observed in influenza virus induced respiratory disease ^31^. Likewise, it will be important to better understand the phenotypic kinetics of SARS-CoV-2 specific T cells and their contribution to long-term protection.

This study has limitations. Foremost is the relatively small sample size. The need to generate a well characterized sample set, limited the number of subjects that could be included. Second, the study is confined to a sampling of COVID-19 convalescent individuals from the greater Baltimore/Washington DC area. As such, this is a geographically restricted population and may not be broadly representative. Third, a low proportion of those who were evaluated had been hospitalized. While this limited our ability to investigate T cell responses in those who were severely ill, it has afforded insight into those with milder disease, which is a more commonly encountered form of COVID-19 and could alternatively be considered a strength of our study. Fourth, while the HLA types which were included account for ~73% of the continental US population, the technology was restricted to only six HLA types. Lastly, the study was cross-sectional and restricted to a relatively narrow time period. Specifically, individuals were evaluated 27-62 days post-symptom resolution. At a minimum, they needed to be at least 28 days post-resolution to donate samples. This limits the conclusions with respect to earlier and/or later in the convalescent period. Of note, even within the period that was evaluated, changes in the T cell and cytokine responses were observed over time. For example, those later in the convalescent period exhibited T cell maturation with effector cells remaining, possibly to clear residual infection. This is consistent with the cytokine data, demonstrating a time effect since diagnosis.

To our knowledge, this is the most comprehensive and precise characterization of SARS-CoV-2-specific CD8+ T cell epitope recognition and corresponding *ex vivo* T cell phenotypes in COVID-19 convalescent subjects to date. The discovery of hitherto undescribed SARS-CoV-2 T cell specificities, their unbiased phenotypic evaluation, and their correlation with the overall inflammation greatly extends the current understanding about natural immunity to SARS-CoV-2. Knowing the combination of epitope targets and T cell profiles capable of differentiating into long-term mediators of protection may be pivotal for triggering a durable immune response. Based on these findings, it seems prudent to include several internal and non-structural viral proteins in the rational design of a second-generation multivalent vaccine.

## Methods

### Sample selection, antibody titers, HLA typing and cytokine testing

The study samples were collected from individuals who were at least 18 years old, who had recovered from COVID-19, and expressed a willingness to donate COVID-19 convalescent plasma (CCP). In order to qualify for CCP donation, individuals had to have a history of COVID-19 as confirmed by a molecular test (e.g. nasopharyngeal swab) for SARS-CoV-2 and meet all eligibility criteria for community blood donation (e.g. not having been pregnant within the six weeks prior to donation, no history or socio-behavioural risk factors for the major transfusion transmissible infections e.g. HIV, hepatitis B and C) ^3^. Eligible individuals were enrolled in the study under full, written informed consent, after which whole blood (25 mL) samples were collected. The samples were separated into plasma and peripheral blood mononuclear cells (PBMC) within 12 hours of blood collection. Aliquots of plasma and PBMC were stored at −80°C until further processing.

A subset of convalescent individuals was selected for evaluating SARS-CoV-2 specific CD8+ T cells using highly multiplexed mass cytometry. Among the first 118 eligible CCP donors, there were 87 individuals with at least four vials of PBMCs collected (each vial contains at least 5 million PBMCs). These individuals were grouped into tertiles (high, medium and low IgG titers) according to overall anti-SARS-CoV-2 IgG titers based on EuroImmun ELISA results against SARS-CoV-2 ^3^ (Table S2). Fifteen individuals were randomly selected from each tertile for HLA typing using the donor PMBC samples. HLA-A and -B loci were tested from genomic DNA by next generation sequencing using the TruSight HLA v1 Sequencing Panel, CareDx^®^, South San Francisco, CA. Individuals matched for ≥2 HLA-A or B alleles (HLA-A*01:01, HLA-A*02:01, HLA-A*03:01, HLA-A*11:01, HLA-A*24:02 and HLA-B*07:02) were included in the subsequent analyses. The remainder of individuals matched for one HLA-A or B allele were randomly selected so that each tertile group comprised ten different donors (total n=30).

The 30 donor samples were transferred to ImmunoScape from JHU in the form of cryopreserved PBMCs. Each sample consisted of either one or two aliquots with an average cell number of 12.15 million cells and a viability above 95% per donor. Samples were thawed at 37°C and immediately transferred into complete RPMI medium (10% hiFCS, 1% penicillin/streptomycin/glutamine, 10mM HEPES, 55μM 2-mercaptoethanol (2-ME) supplemented with 50 U/ml Benzonase (Sigma). Aliquots derived from the same donors were combined and all samples were enriched for T cells by removing CD14 and CD19 expressing cells using a column-based magnetic depletion approach according to the manufacturer’s recommendations (Miltenyi). Healthy donor PBMCs (STEMCELL) matched for at least one of the donor HLA alleles were included in each experiment as control for specific T cell identification.

SARS-CoV-2 neutralizing antibody (nAbs) titers against 100 50% tissue culture infectious doses (TCID_50_) per 100 uL were determined using a microneutralization (NT) assay, as previously described ^3^. The nAb titer was calculated as the highest plasma dilution that prevented cytopathic effect (CPE) in 50% of the wells tested. nAb area under the curve (AUC) values were estimated using the exact number of wells protected from infection at every plasma dilution.

Highly sensitive, multiplexed sandwich immunoassays using MULTI-ARRAY^®^ electrochemiluminescence detection technology (MesoScale Discovery, Gaithersburg, MD, USA) were used for the quantitative evaluation of 35 different human cytokines and chemokines in plasma samples from eligible CCP donors [IFN-γ, IL-1β, IL-2, IL-4, IL-6, IL-8, IL-10, IL-12p70, IL-13, TNF-α, GM-CSF, IL-1α, IL-5, IL-7, IL-12/IL23p40, IL-15, IL-16, IL-17A, TNF-β, VEGF-A, Eotaxin, MIP-1β, Eotaxin-3, TARC, IP-10, MIP-1α, MCP-1, MDC, MCP-4, IL-18, IL-1RA, G-CSF (CSF3), IFN-α2a, IL-33 and IL-21]. Cytokine and chemokine concentrations were calculated per manufacturer protocol (MSD DISCOVERY WORKBENCH^®^ analysis software) and were considered “detectable” if both runs of each sample had a signal greater than the analyte- and plate-specific lower limit of detection (LLOD) (i.e., 2.5 standard deviations of the plate-specific blank). Cytokine and chemokine concentrations (pg/mL) from both runs of each analyte were averaged.

### Peptides

A total of 408 unique SARS-CoV2 candidate peptide epitopes spanning six HLAs (HLA-A*01:01, HLA-A*02:01, HLA-A*03:01, HLA-A*11:01, HLA-A*24:02 and HLA-B*07:02) were selected based on recent predictions ^14,15^ (Table S3). For each of the HLA alleles tested, up to 20 different control peptides (SARS-CoV-2 unrelated epitopes) were also included into the screenings (Table S3). All peptides were ordered from Genscript (China) or Mimotopes, (Australia) with a purity above 85% by HPLC purification and mass spectrometry. Lyophilized peptides were reconstituted at a stock concentration of 10 mM in DMSO.

### Antibody staining panel setup

Purified antibodies lacking carrier proteins (100 μg/antibody) were conjugated to DN3 MAXPAR chelating polymers loaded with heavy metal isotopes following the recommended labelling procedure (Fluidigm). A specific staining panel was set up consisting of 28 antibodies addressing lineage, phenotypic and functional markers (Table S5). All labelled antibodies were titrated and tested by assessing relative marker expression intensities on relevant immune cell subsets in PBMCs from healthy donors (STEMCELL). Antibody mixtures were prepared freshly and filtered using a 0.1 mM filter (Millipore) before staining.

### Tetramer multiplexing setup

To screen for SARS-CoV-2-specific CD8+ T cells we set up a three-metal combinatorial tetramer staining approach as described previously ^17,32^. Briefly, specific peptide-MHC class I complexes were generated by incubating biotinylated UV-cleavable peptide HLA monomers in the presence of individual antigen candidates. For the generation of a triple-coded tetramer staining mixture recombinant streptavidin was conjugated to heavy metal loaded DN3 polymers ^17^ and three out of 12 differently labelled streptavidin molecules were randomly combined by using an automated pipetting device (TECAN) resulting in a total of 220 unique possible combinations to encode single peptide candidates. Peptide exchange was performed at 100μg/mL of HLA monomer in PBS with 50μM peptides of interest in a 96-well plate. Peptides with similar sequences were assigned the same triple code to avoid multiple signals through potential T cell cross-reactivity. According to the donors’ HLA genotypes, total epitope screenings ranged from 49 to 220 peptides for individual samples, including SARS-CoV-2 unrelated control peptides. For tetramerization, each triple coded streptavidin mixture was added in three steps to their corresponding exchanged peptide– MHC complexes to reach a final molar ratio of 1:4 (total streptavidin:peptide–MHC). The tetramerized peptide–MHC complexes were incubated with 10μM of free biotin (SIGMA) to saturate remaining unbound streptavidins. All tetramers were combined and concentrated (10 kDa cutoff filter) in cytometry buffer (PBS, 2% fetal calf serum, 2 mM EDTA, 0.05% sodium azide) before staining the cells. As internal control and to facilitate the detection of *bona fide* antigen-specific T cells we generated a second tetramer staining configuration for each experiment using a completely different coding scheme for each peptide ^17^.

### Sample staining and acquisition

T cell enriched donor samples and healthy donor PBMCs were split into two fractions and seeded at equal numbers in two wells of a 96 well plate. Cells were washed and each well was then stained with 100ul of either one of the two tetramer configurations for 1h at RT. After 30 mins a unique metal (Cd-111 and Cd-113) labelled anti-CD45 antibody was added into each of the wells to further barcode the cells that were stained with the different tetramer configurations. Cells were then washed twice and the two wells per sample were combined and stained with the heavy metal labelled antibody mixtures for 30 mins on ice and 200 μM cisplatin during the last 5 mins for the discrimination of live and dead cells. Cells were washed and fixed in 2% paraformaldehyde in PBS overnight at 4°C. For intracellular staining, cells were incubated in 1× permeabilization buffer (Biolegend) for 5 min on ice and incubated with metal conjugated anti-GranzymeB antibodies for 30 min on ice. Samples from different donors were barcoded with a unique dual combination of bromoacetamidobenzyl-EDTA (Dojindo)-linked metal barcodes (Pd-102, Pd-104, PD106 and PD108, and Pd-110) for 30 min on ice. Cells were then washed and resuspended in 250 nM iridium DNA intercalator (Fuidigm) in 2% paraformaldehyde/PBS at RT. Cells were washed, pooled together and adjusted to 0.5 million cells per ml H_2_O together with 1% equilibration beads (EQ Four element calibration beads, Fluidigm) for acquisition on a HELIOS mass cytometer (CyTOF, Fluidigm).

### Data analysis

After mass cytometry acquisition, signals for each parameter were normalized based on EQ beads (Fluidigm) added to each sample ^33^ and any zero values were randomized using a custom Rscript that uniformly distributes values between minus-one and zero. Each sample was manually de-barcoded followed by gating on live CD8+ and CD4+ T cells (CD45^+^ DNA^+^ cisplatin- CD3^+^ cells) from either staining configuration after gating out residual monocytes (CD14) and B cells (CD19) using FlowJo (Tree Star Inc) software. Antigen-specific triple tetramer positive cells (hits) were identified by an automated peptide-MHC gating method ^17^ and each hit was confirmed and refined using manual gating. The designation of *bona fide* antigen-specific T cells was further dependent on (i) the detection cut-off threshold (≥2 events to be detected in each staining configuration), (ii) the frequency correspondence between *the two tetramer staining configurations (*ratio between the frequencies of a hit in either staining configuration to be ≤2) and (iii) the background noise (frequencies of specific CD8 T+ cell events must be greater than events from the corresponding CD4 T cell population), as unbiased objective criteria for antigen-specificity assessment ^16^. Bulk T cells and true hits from both staining configurations were combined for assessing frequencies, phenotypic and statistical analysis.

Frequency values were calculated based on the percentage of the parent immune cell population. Phenotypic markers were gated individually for each sample and calculated as % of positive cells. High-dimensional phenotypic profiles and sample distributions were shown using uniform manifold approximation and projection ^34^ and Phenograph for automated cell clustering ^35^. Data analysis was performed using CYTOGRAPHER®, immunosCAPE’s cloud based analytical software, custom R-scripts, GraphPad Prism and Flowjo software.

### Statistical analysis

Comparative analyses of frequencies of cell subsets and marker expression between samples were done using Wilcoxon rank sum tests, extended to Kruskall-Wallis tests by ranks for more than 2 levels in a grouping variable; resulting p-values were adjusted for multiple testing using the Benjamini-Hochberg method to control the false discovery rate. Correlations were calculated with the Spearman’s rank-order test. A correlation matrix was calculated comparing phenotypic and serological marker variables in a pairwise fashion, using the *corr.test* function from the *psych* CRAN package; the *corrplot* package was subsequently used to graphically display the correlation matrix. Resulting p-values were adjusted for multiple testing using the Bonferroni method. Spearman’s correlation coefficients were indicated by a heat scale whereby blue color shows positive linear correlation, and red color shows negative linear correlation. All statistical analyses were performed using GraphPad Prism and R and statistical significance was set at a threshold of *p < 0.05, **p < 0.01, and ***p < 0.001.

## Acknowledgements

The authors would like to thank the CCP study and laboratory teams, and all the donors for the generous participation in the study. We thank the National Institute of Infectious Diseases, Japan, for providing VeroE6TMPRSS2 cells and acknowledge the Centers for Disease Control and Prevention, BEI Resources, NIAID, NIH for SARS-Related Coronavirus 2, Isolate USA-WA1/2020, NR-5228.

## Contributions

Contributed samples – E.B., A.T., D.S., S.S.

Collected experimental data – T.B., K.L, A.P, H.S., F.K., M.B. and M.F.

Analysed data and performed statistical analysis H.K. D.C., E.P. and M.F.

Drafted the manuscript H.K., B.A.,E.W.N., A.N. and M.F.

Conceived and designed the study: B.A., E.W.N., A.N., MF., A.T., A.R., E.B. and T.Q.

All authors contributed intellectually and approved the manuscript

---

**Supplementary Figure 1.**
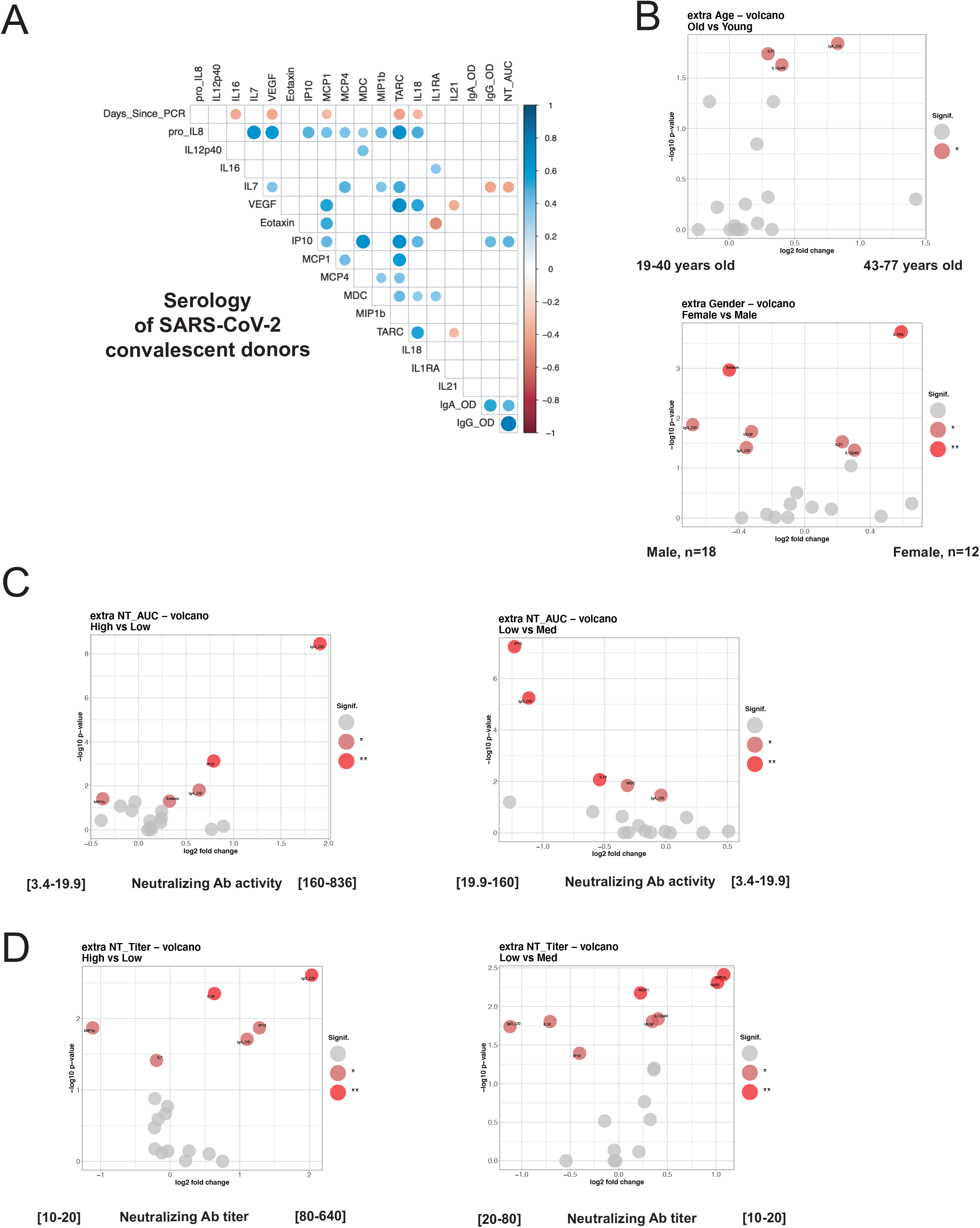
Correlations between antibody titers, cytokines, neutralizing antibody activity and recovery time in convalescent donor cross-sectional sample

**Supplementary Figure 2.**
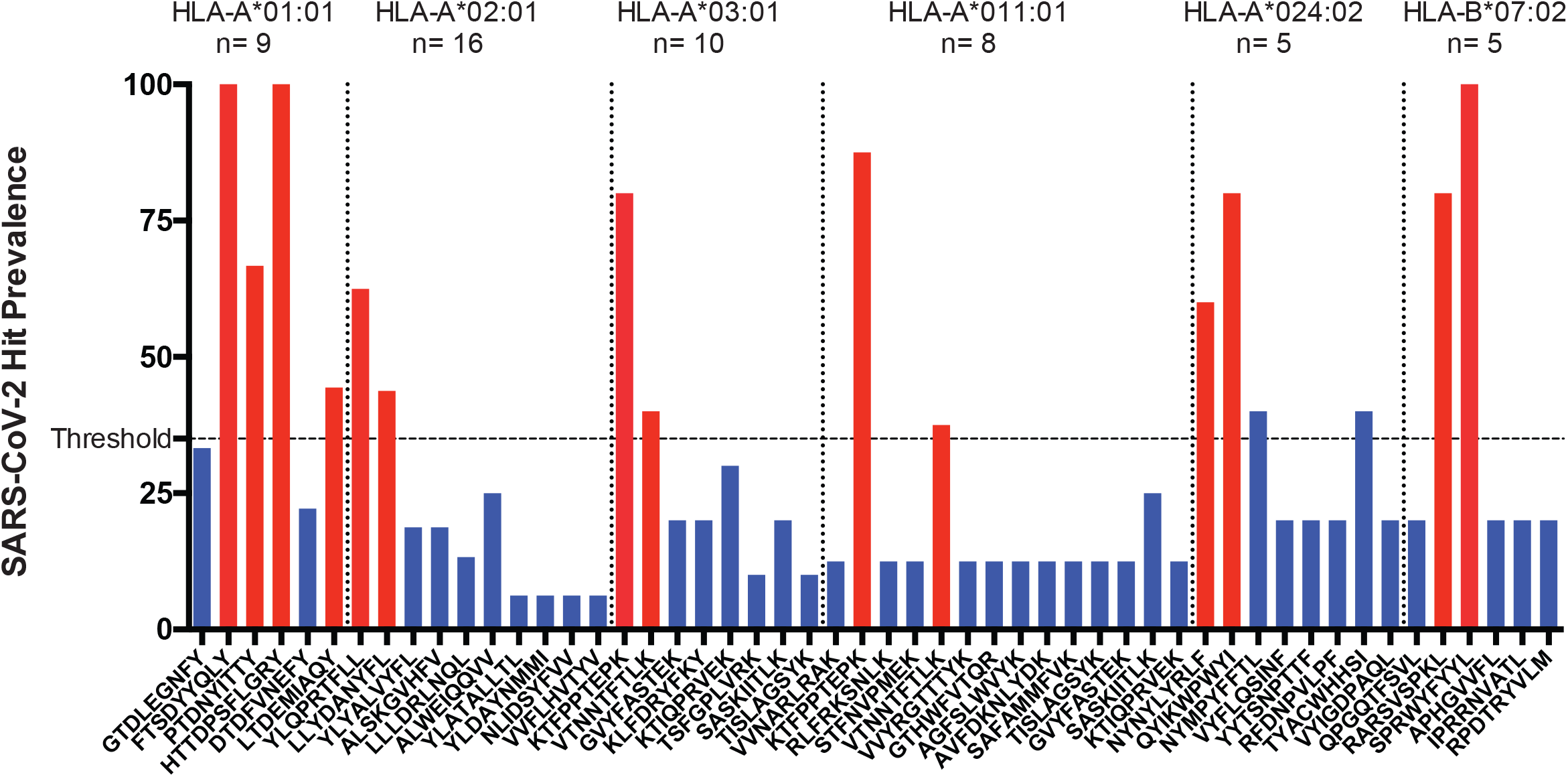
Definition of SARS-CoV-2 hit prevalence responses across all HLAs

**Supplementary Figure 3.**
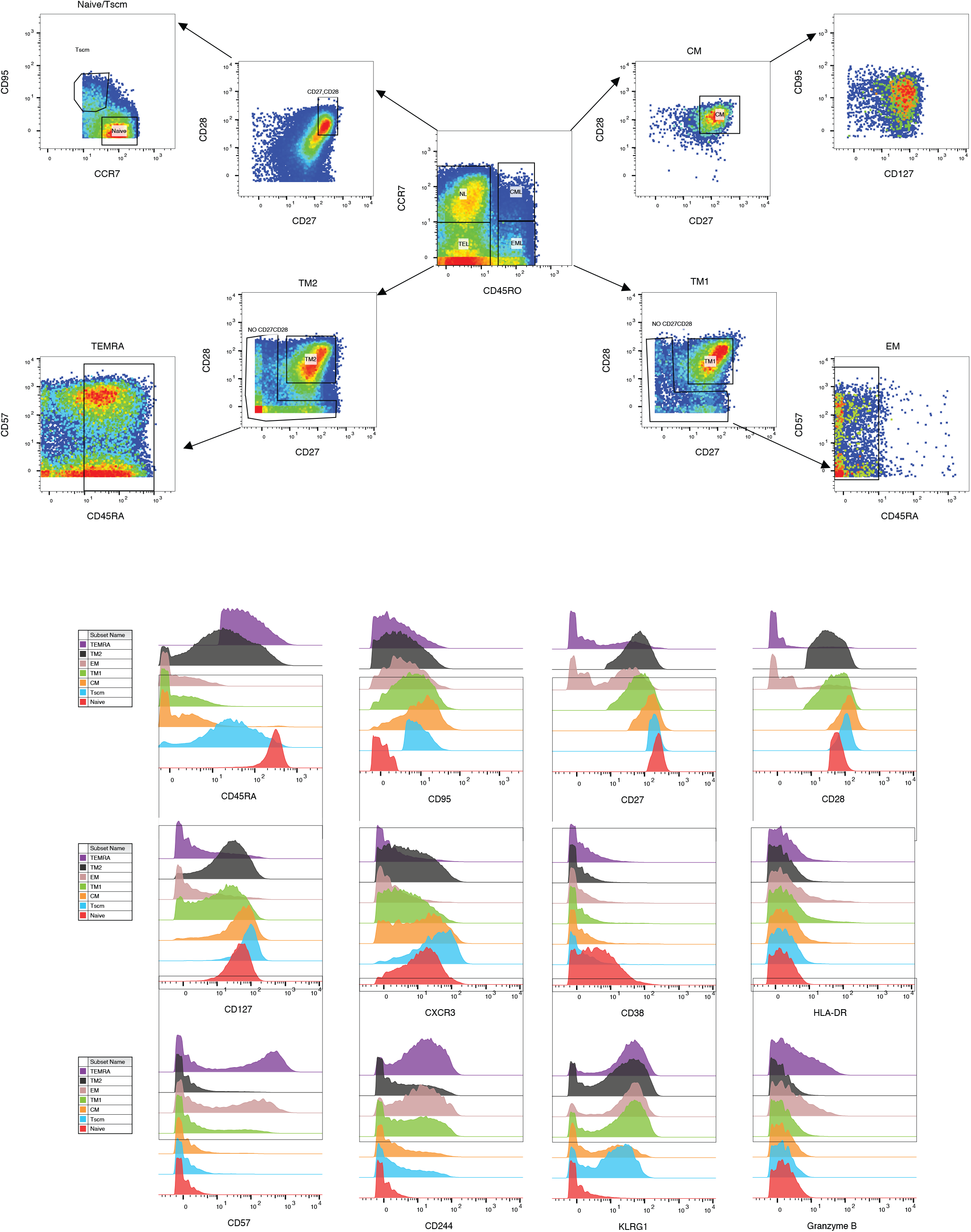
Gating scheme for identification of T cell differentiation states

**Supplementary Figure 4.**
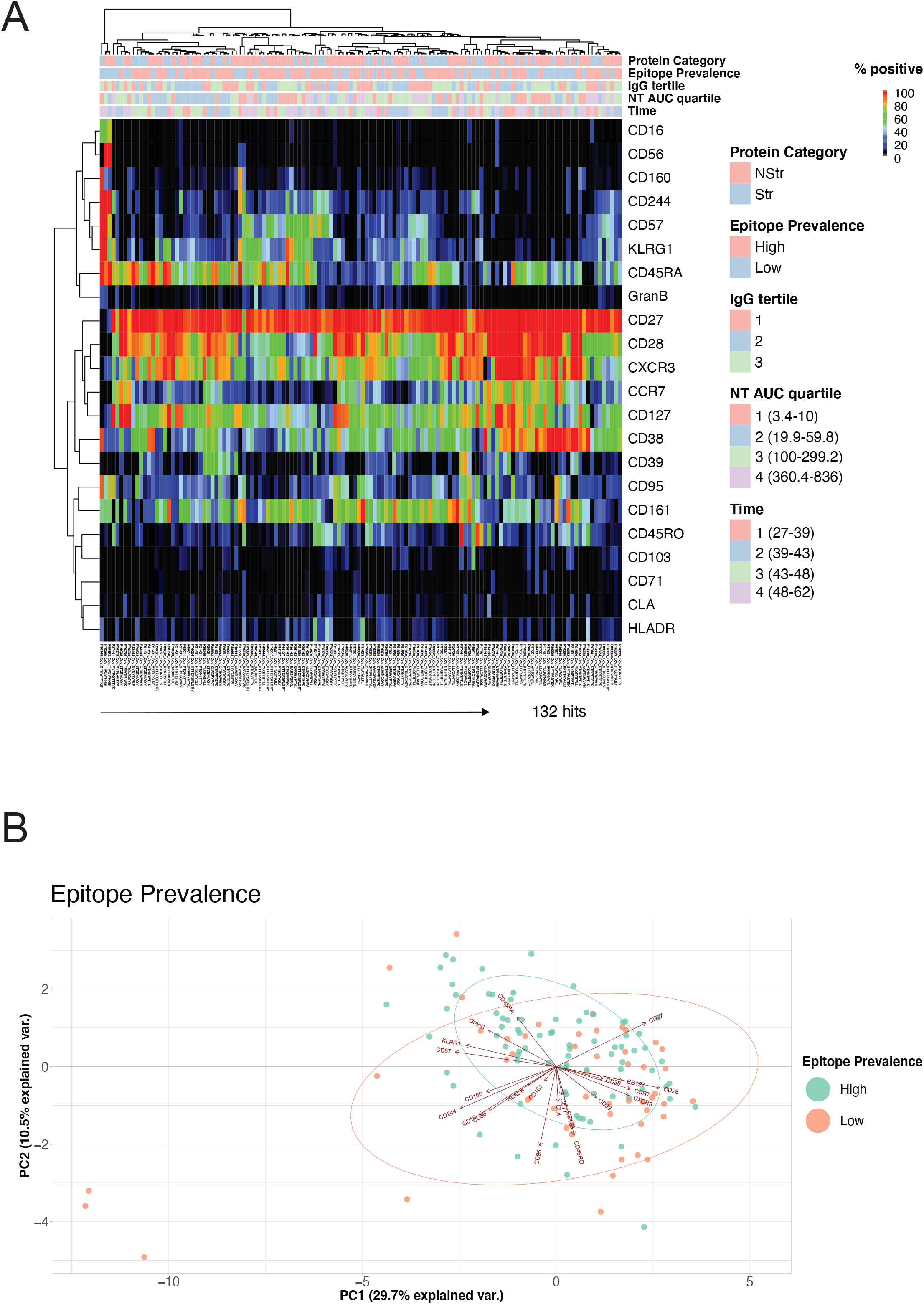
Association of SARS-CoV-2-specific CD8+ T cells with clinical parameters and epitope categories

**Supplementary Figure 5.**
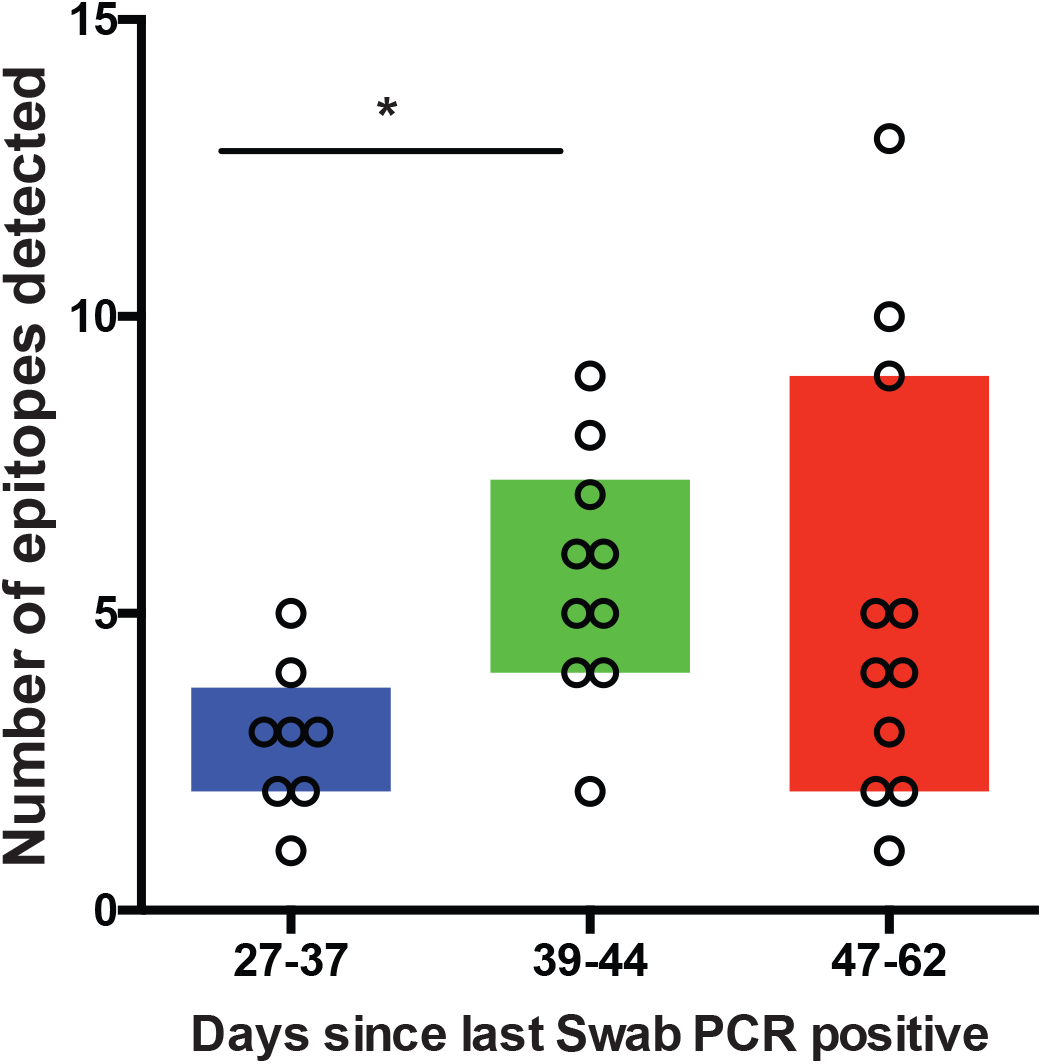
Numbers of epitopes detected over time

**Supplementary Figure 6.**
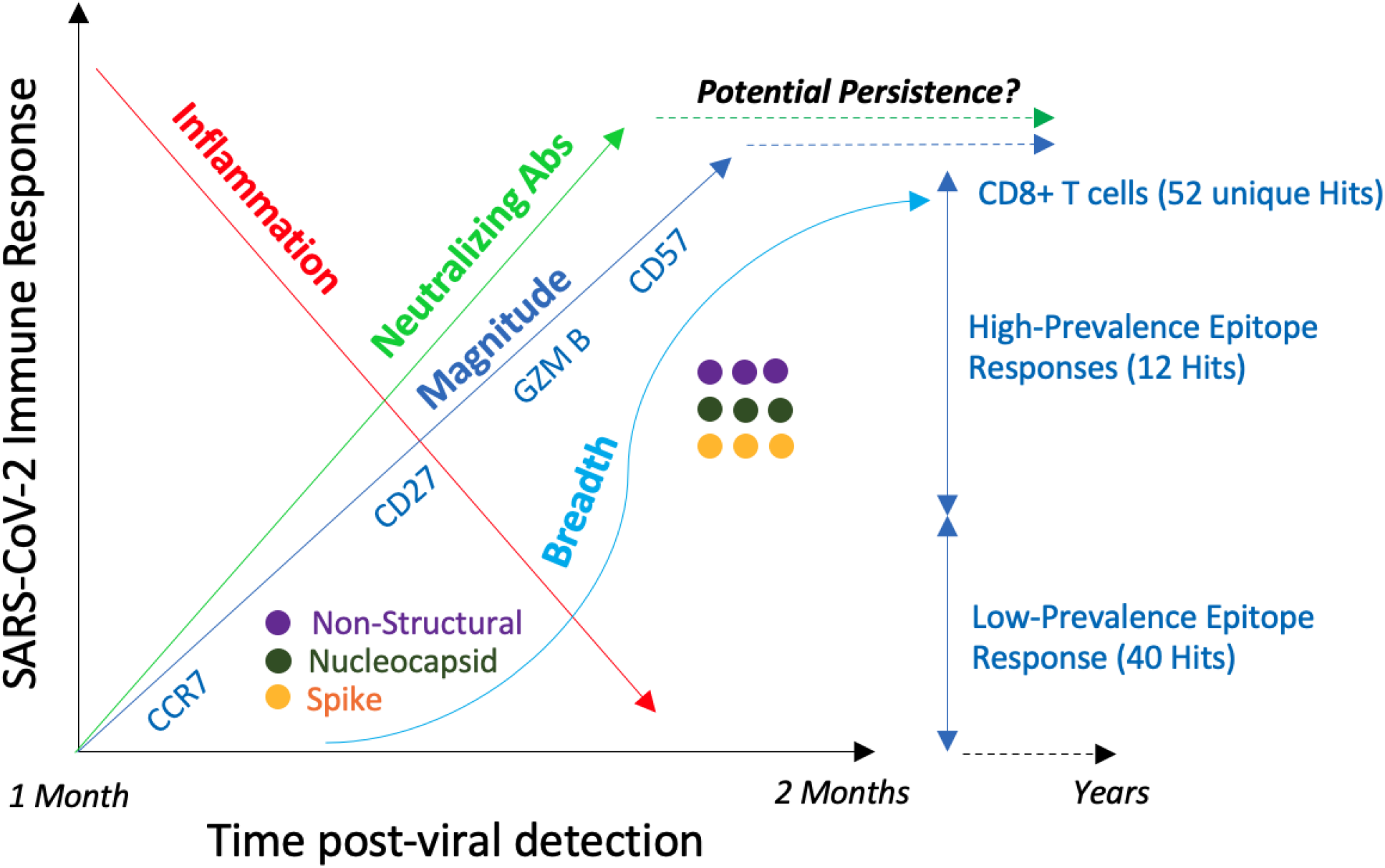
Summary model

**Table S1.**

| Sample | CCP ID | Race | Gender | Age | Hospitalization | Days since swab | IgG OD (pos >0.8) | IgA OD (pos >4) | IgG Avididity IC50 | NTAUC | NTtiter |
| --- | --- | --- | --- | --- | --- | --- | --- | --- | --- | --- | --- |
| 1 | C14870 | Hispanic | Male | 53 | no | 48 | 4.5 | 2.7 | 4.1 | 618.4 | 320 |
| 2 | C16009 | white | Male | 32 | no | 41 | 1.9 | 4.2 | 4 | 139.6 | 80 |
| 3 | C17767 | white | Male | 28 | no | 37 | 1.4 | 2.4 | 4 | 10 | 20 |
| 4 | C20061 | white | Female | 62 | no | 42 | 6.8 | 3.6 | 4 | 30.1 | 40 |
| 5 | C20750 | mixed/other/unknown | Female | 68 | no | 30 | 3.7 | 3.1 | 3.9 | 100 | 80 |
| 6 | C21431 | white | Male | 29 | no | 43 | 4.3 | 1.8 | 4.6 | 19.9 | 20 |
| 7 | C31691 | white | Female | 61 | no | 27 | 0.4 | 0.4 | 3.8 | 10 | 20 |
| 8 | C36143 | white | Female | 77 | no | 51 | 1.9 | 8.2 | 3.1 | 10 | 20 |
| 9 | C36346 | white | Male | 56 | no | 37 | 4.5 | 3.1 | 3.8 | 54.7 | 40 |
| 10 | C37228 | white | Female | 50 | no | 33 | 0.1 | 0.1 | N.A. | 3.4 | 10 |
| 11 | C37621 | white | Male | 43 | no | 41 | 1.4 | 1.9 | 3.7 | 3.4 | 10 |
| 12 | C43444 | white | Male | 19 | no | 43 | 4.7 | 2.1 | 3.9 | 169 | 160 |
| 13 | C44137 | white | Female | 31 | no | 47 | 0.6 | 0.3 | 4.3 | 10 | 20 |
| 14 | C45611 | white | Male | 40 | no | 37 | 2.8 | 3.2 | 3.6 | 30.1 | 40 |
| 15 | C48630 | white | Male | 50 | yes (1 day) | 47 | 10.5 | 6 | 5.1 | 541.6 | 320 |
| 16 | C51850 | white | Female | 61 | no | 42 | 3.7 | 2.5 | 4.4 | 139.6 | 80 |
| 17 | C63869 | white | Male | 32 | no | 57 | 2.8 | 0.8 | 4.2 | 3.4 | 10 |
| 18 | C70293 | Asian | Female | 43 | no | 50 | 1.8 | 1.7 | 3.5 | 6.7 | 20 |
| 19 | C70295 | white | Male | 37 | no | 48 | 7.2 | 3 | 5.1 | 240.4 | 160 |
| 20 | C74517 | white | Male | 28 | no | 50 | 2.3 | 0.9 | 4.6 | 59.8 | 40 |
| 21 | C74999 | Asian | Male | 48 | no | 35 | 7.5 | 9.4 | 5.5 | 541.6 | 320 |
| 22 | C76378 | white | Male | 64 | no | 40 | 3.2 | 1.6 | 3.9 | 160 | 80 |
| 23 | C76624 | Asian | Female | 19 | no | 44 | 2.4 | 0.5 | 4.1 | 19.9 | 20 |
| 24 | C89395 | white | Male | 40 | no | 52 | 5.4 | 3.7 | 4.5 | 460 | 320 |
| 25 | C89892 | white | Male | 37 | no | 39 | 1.9 | 3 | 3.9 | 19.9 | 20 |
| 26 | C91439 | white | Male | 63 | no | 47 | 6.7 | 8.1 | 3.9 | 836 | 640 |
| 27 | C93871 | mixed/other/unknown | Female | 27 | N.A. | 39 | 0.8 | 1 | 4.1 | 59.8 | 40 |
| 28 | C97441 | white | Female | 40 | yes (5days) | 62 | 10.2 | 2.4 | 7.7 | 360.4 | 320 |
| 29 | C97651 | white | Female | 28 | no | 53 | 3.1 | 1.7 | 4.1 | 10 | 20 |
| 30 | C98638 | white | Male | 38 | no | 32 | 4.2 | 1.8 | 3.4 | 299.2 | 160 |

**Table S2.**

| Sample | CCP Number: | IgG tertile | HLAA01 | HLA A02 | HLA A03 | HLA A11 | HLA A24 | HLA B07 |
| --- | --- | --- | --- | --- | --- | --- | --- | --- |
| 1 | C20750 | 2 |  | HLA A02 |  | HLA A11 |  |  |
| 2 | C17767 | 3 |  | HLA A02 | HLA A03 |  |  |  |
| 3 | C16009 | 3 | HLAA01 |  |  |  |  |  |
| 4 | C45611 | 2 | HLAA01 |  | HLA A03 |  |  | HLA B07 |
| 5 | C76378 | 2 |  | HLA A02 |  |  |  | HLA B07 |
| 6 | C31691 | 3 |  |  | HLA A03 |  |  |  |
| 7 | C37228 | 3 |  | HLA A02 |  | HLA A11 |  |  |
| 8 | C98638 | 2 |  |  | HLA A03 |  |  |  |
| 9 | C43444 | 1 |  | HLA A02 |  |  | HLA A24 |  |
| 10 | C74517 | 2 | HLAA01 | HLA A02 |  |  |  |  |
| 11 | C89395 | 1 | HLAA01 |  |  |  |  |  |
| 12 | C93871 | 3 |  | HLA A02 |  | HLA A11 |  |  |
| 13 | C91439 | 1 |  | HLA A02 |  |  |  |  |
| 14 | C51850 | 2 | HLAA01 | HLA A02 |  |  |  |  |
| 15 | C70293 | 3 |  |  | HLA A03 | HLA A11 |  |  |
| 16 | C14870 | 1 |  | HLA A02 |  |  |  |  |
| 17 | C37621 | 3 |  | HLA A02 | HLA A03 |  |  | HLA B07 |
| 18 | C74999 | 1 |  |  |  | HLA A11 | HLA A24 |  |
| 19 | C76624 | 2 | HLAA01 |  |  | HLA A11 |  |  |
| 20 | C70295 | 1 |  |  | HLA A03 |  | HLA A24 | HLA B07 |
| 21 | C97651 | 2 |  | HLA A02 |  |  |  |  |
| 22 | C21431 | 2 | HLAA01 |  |  |  | HLA A24 |  |
| 23 | C89892 | 3 |  |  | HLA A03 |  | HLA A24 |  |
| 24 | C36143 | 3 |  |  | HLA A03 | HLA A11 |  | HLA B07 |
| 25 | C36346 | 1 |  | HLA A02 |  |  |  |  |
| 26 | C44137 | 3 | HLAA01 | HLA A02 |  |  |  |  |
| 27 | C48630 | 1 |  | HLA A02 |  |  |  |  |
| 28 | C63869 | 2 |  | HLA A02 |  |  |  |  |
| 29 | C97441 | 1 |  |  |  | HLA A11 |  |  |
| 30 | C20061 | 1 | HLAA01 |  | HLA A03 |  |  |  |

**Table S3.**
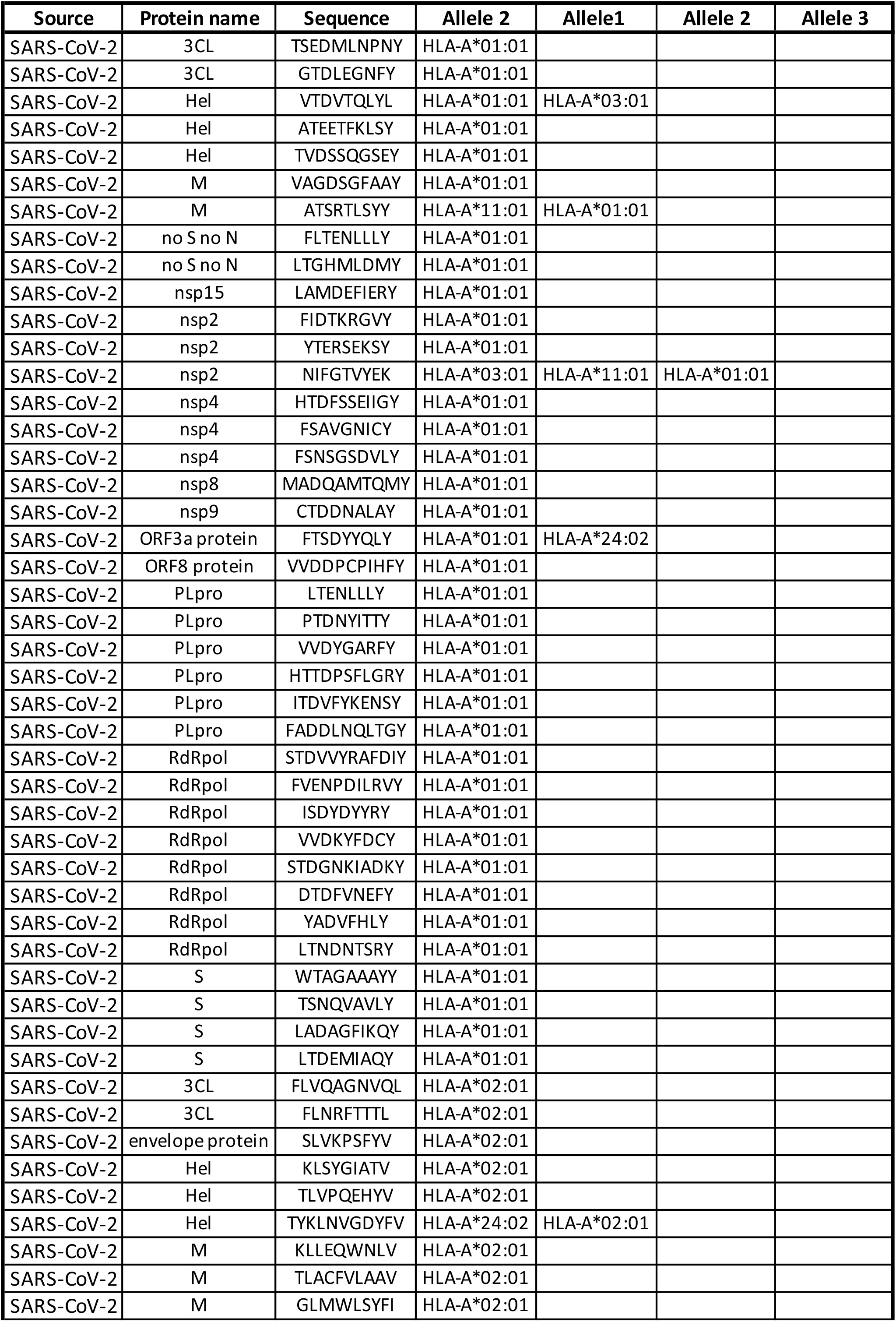
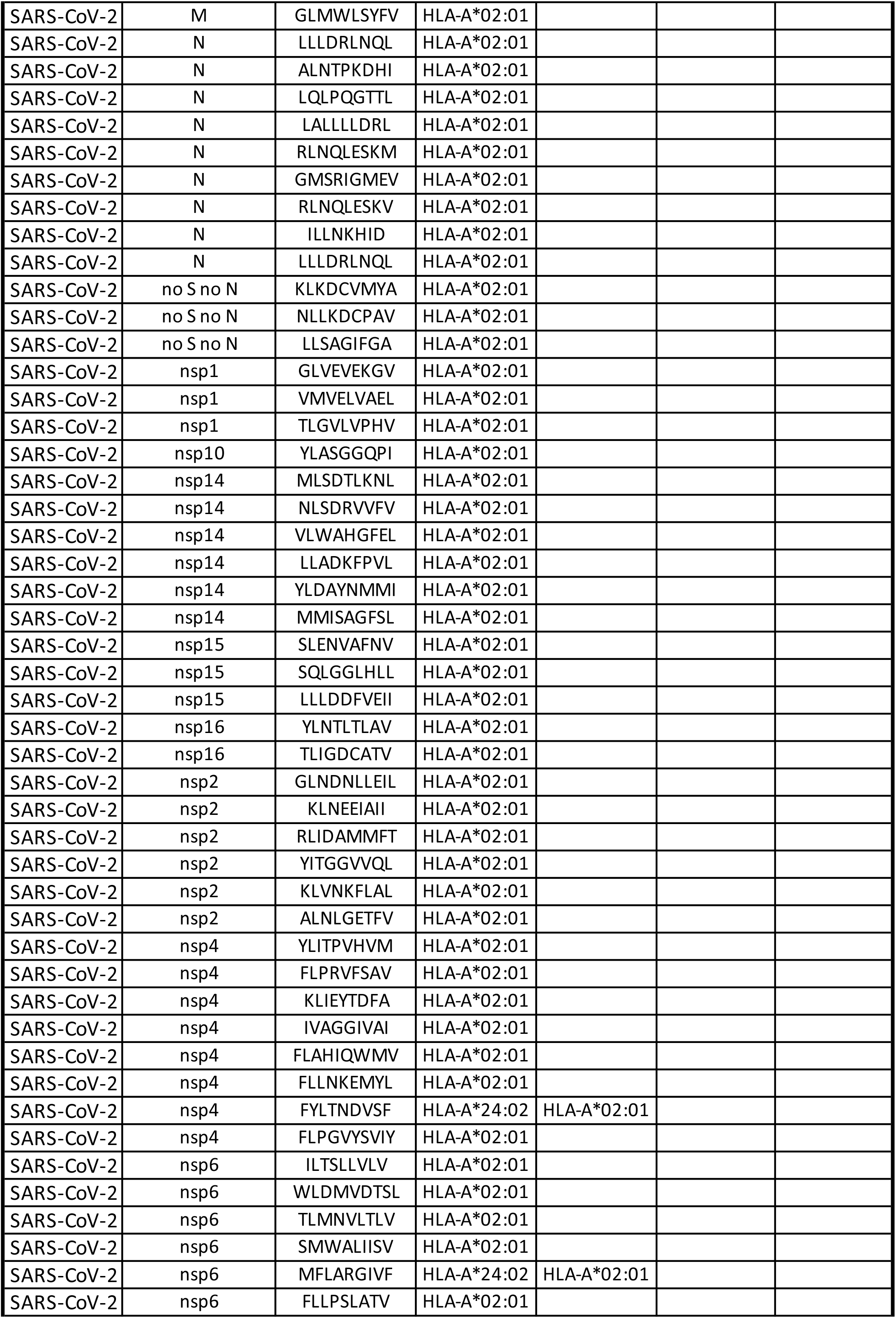
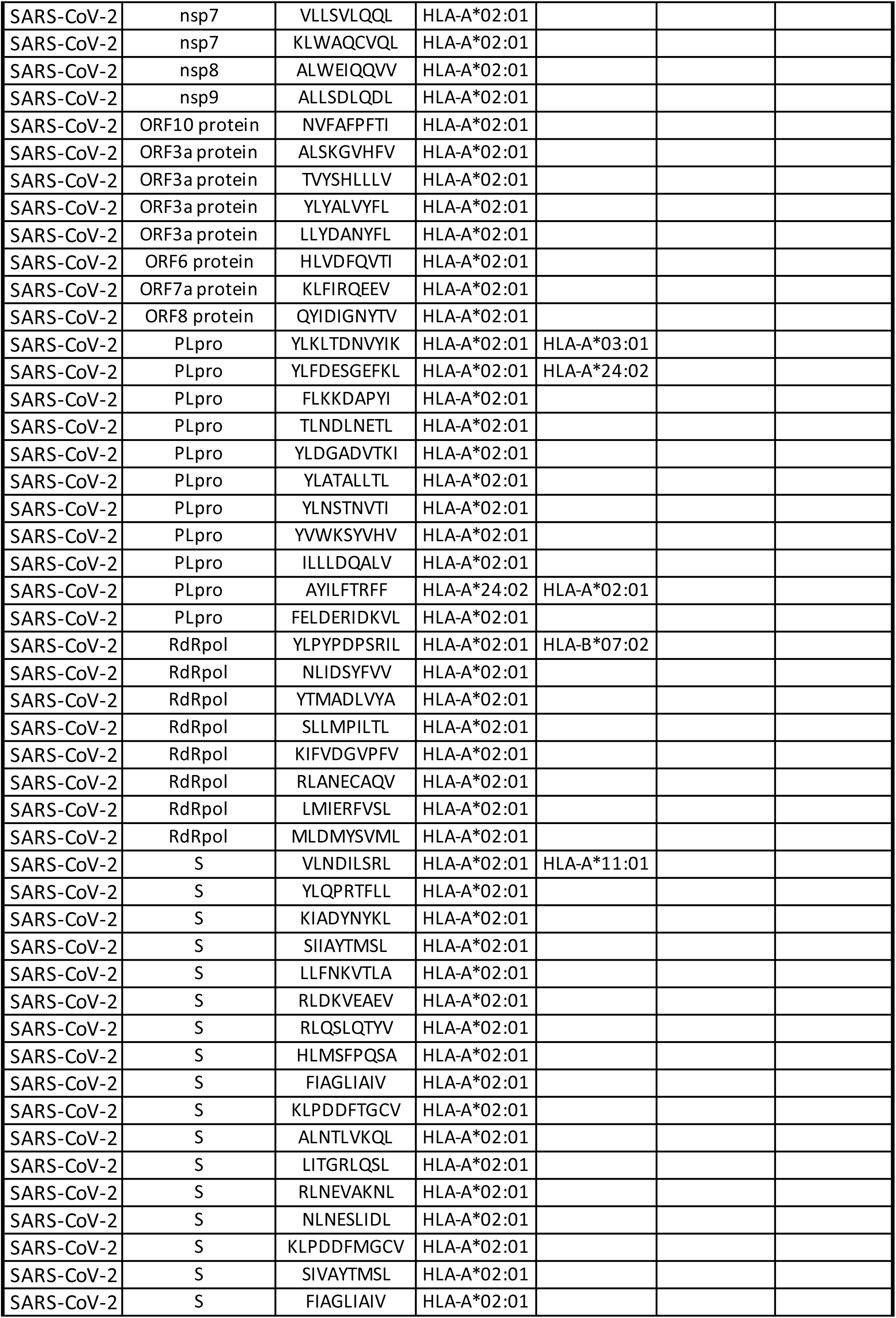
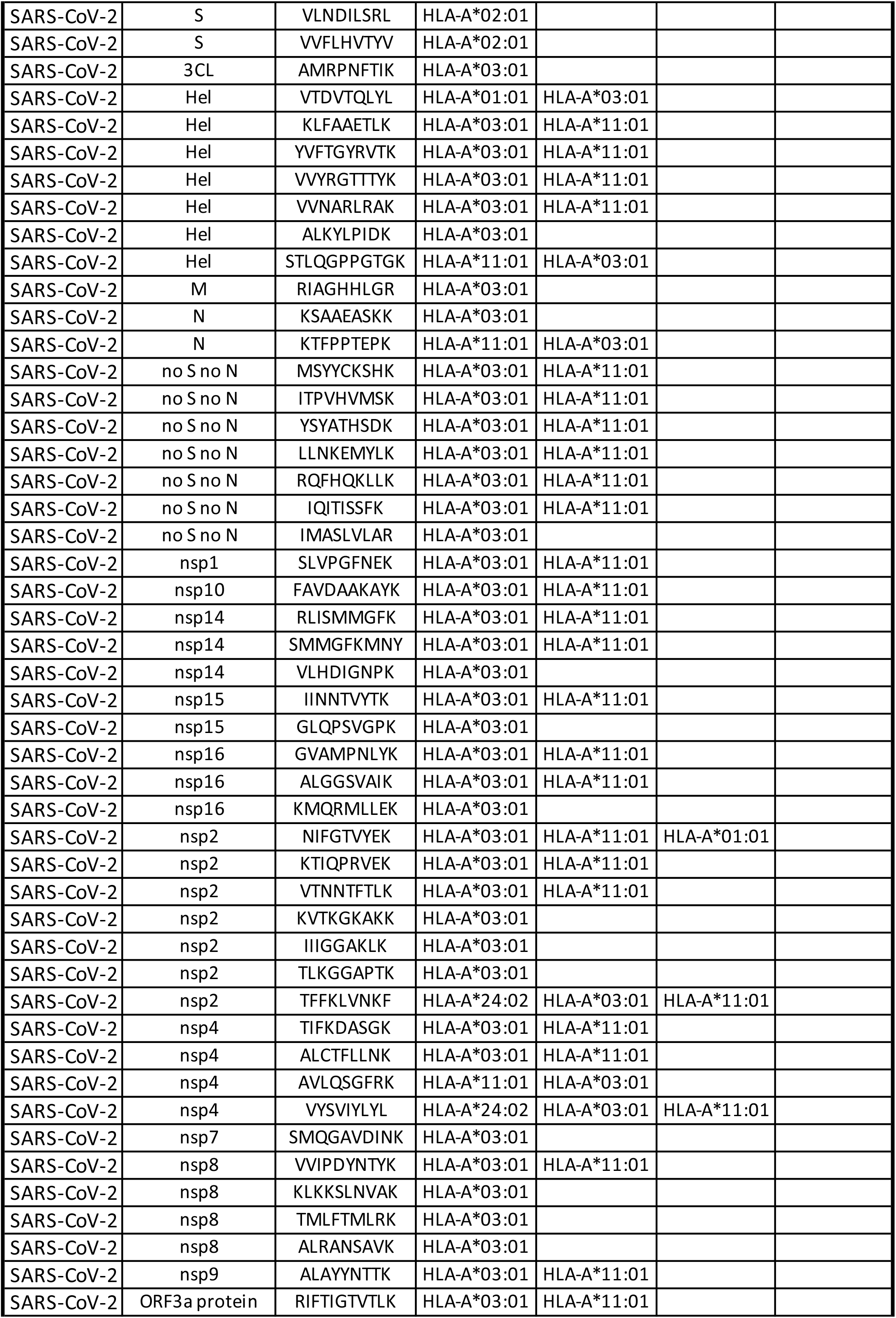
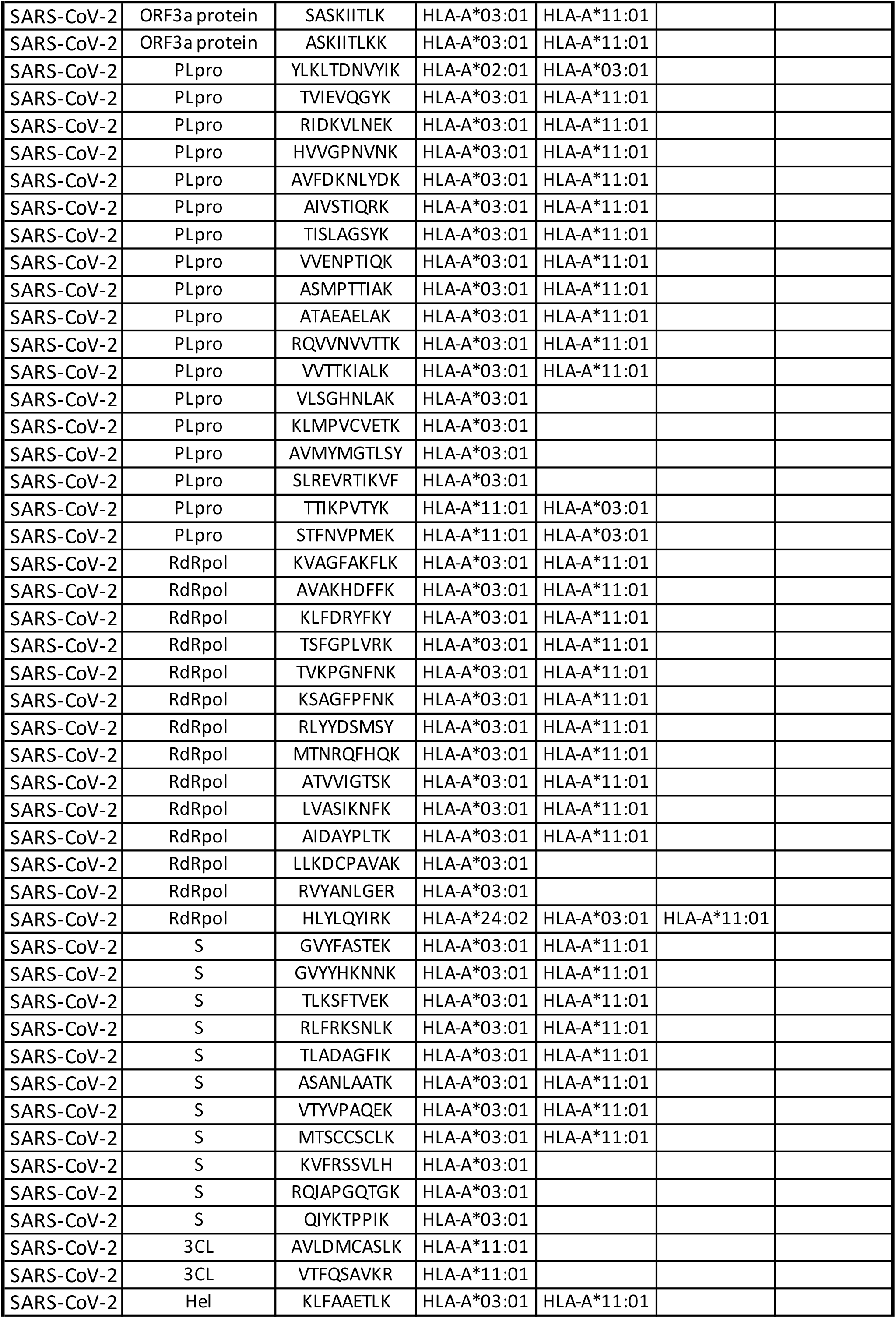
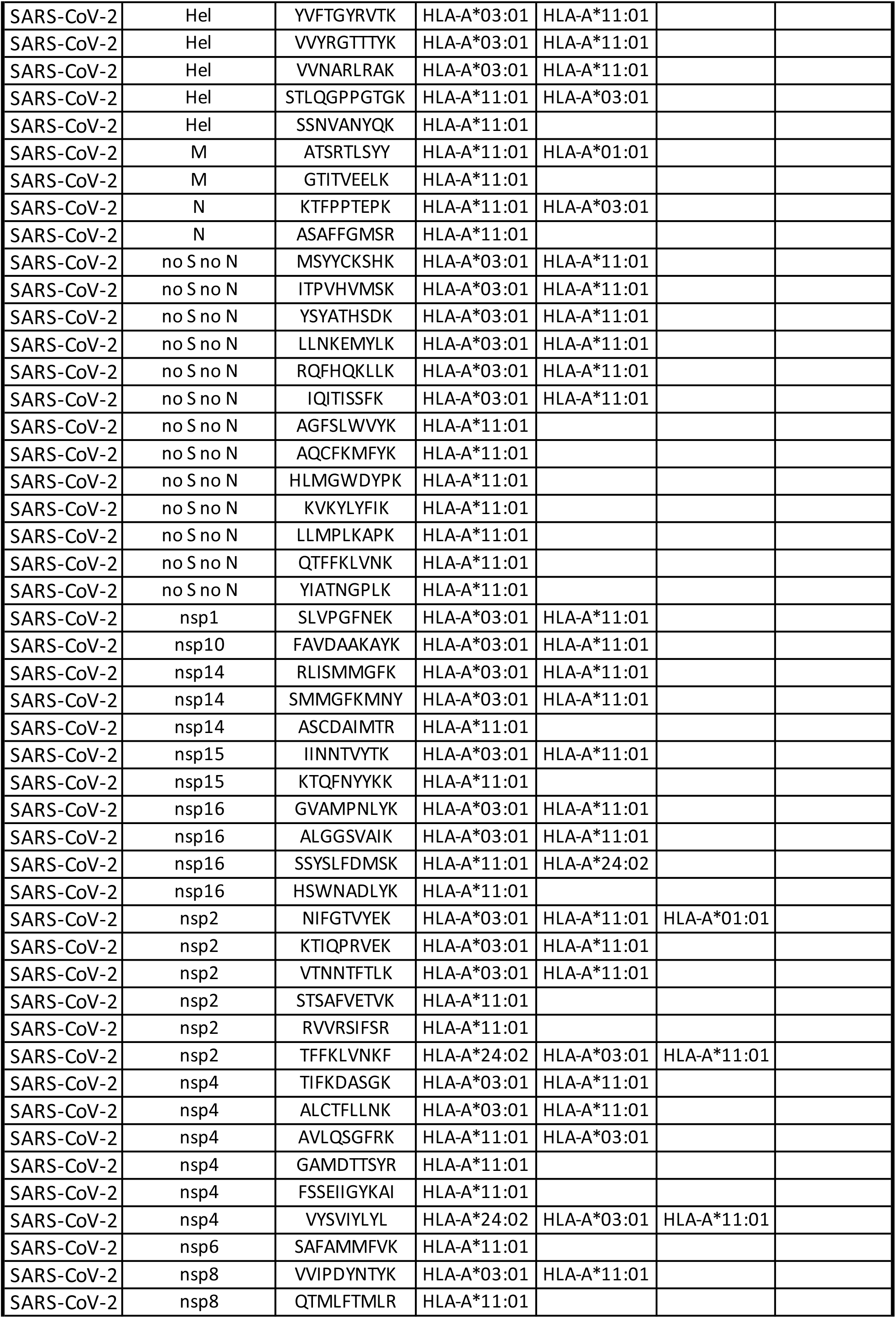
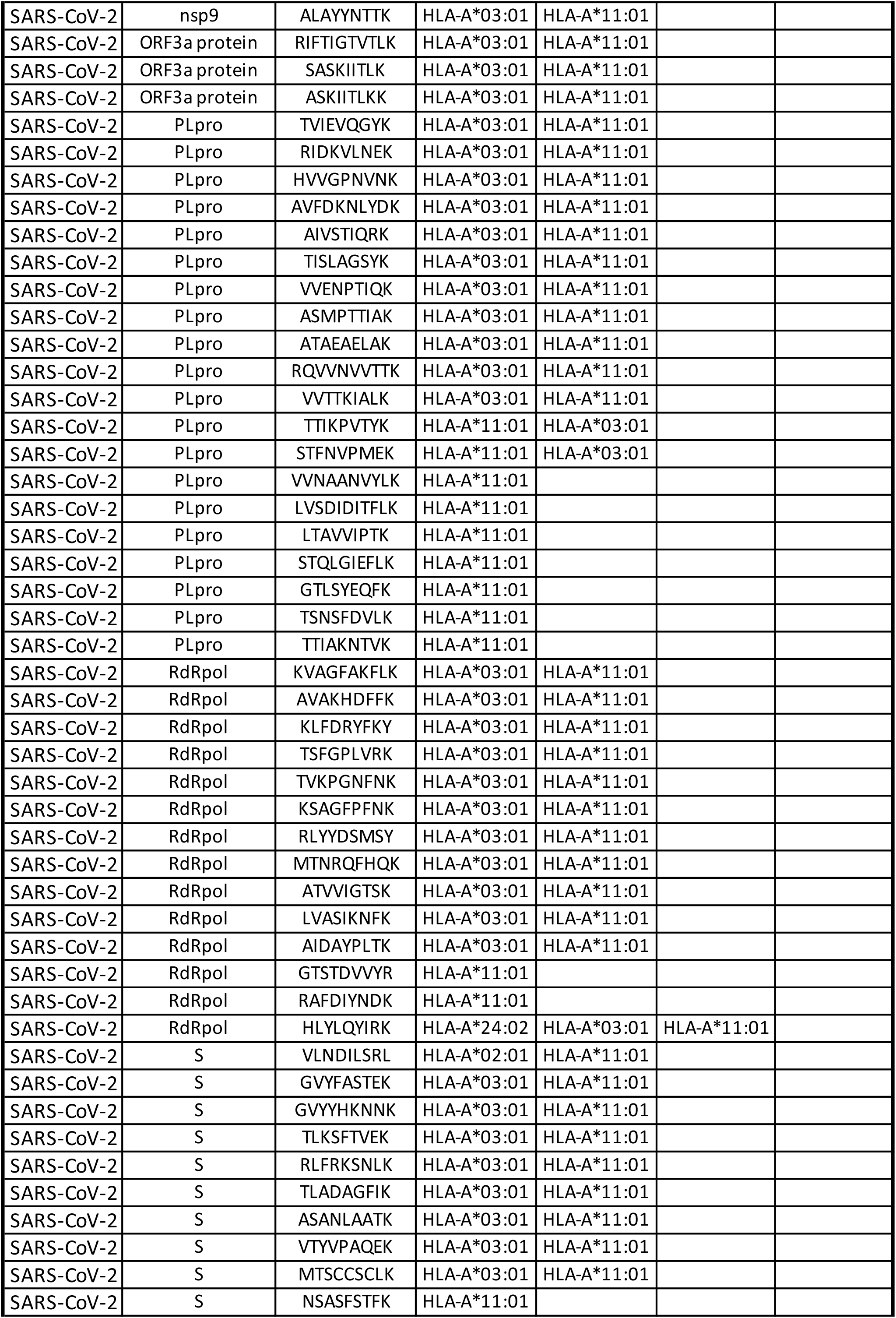
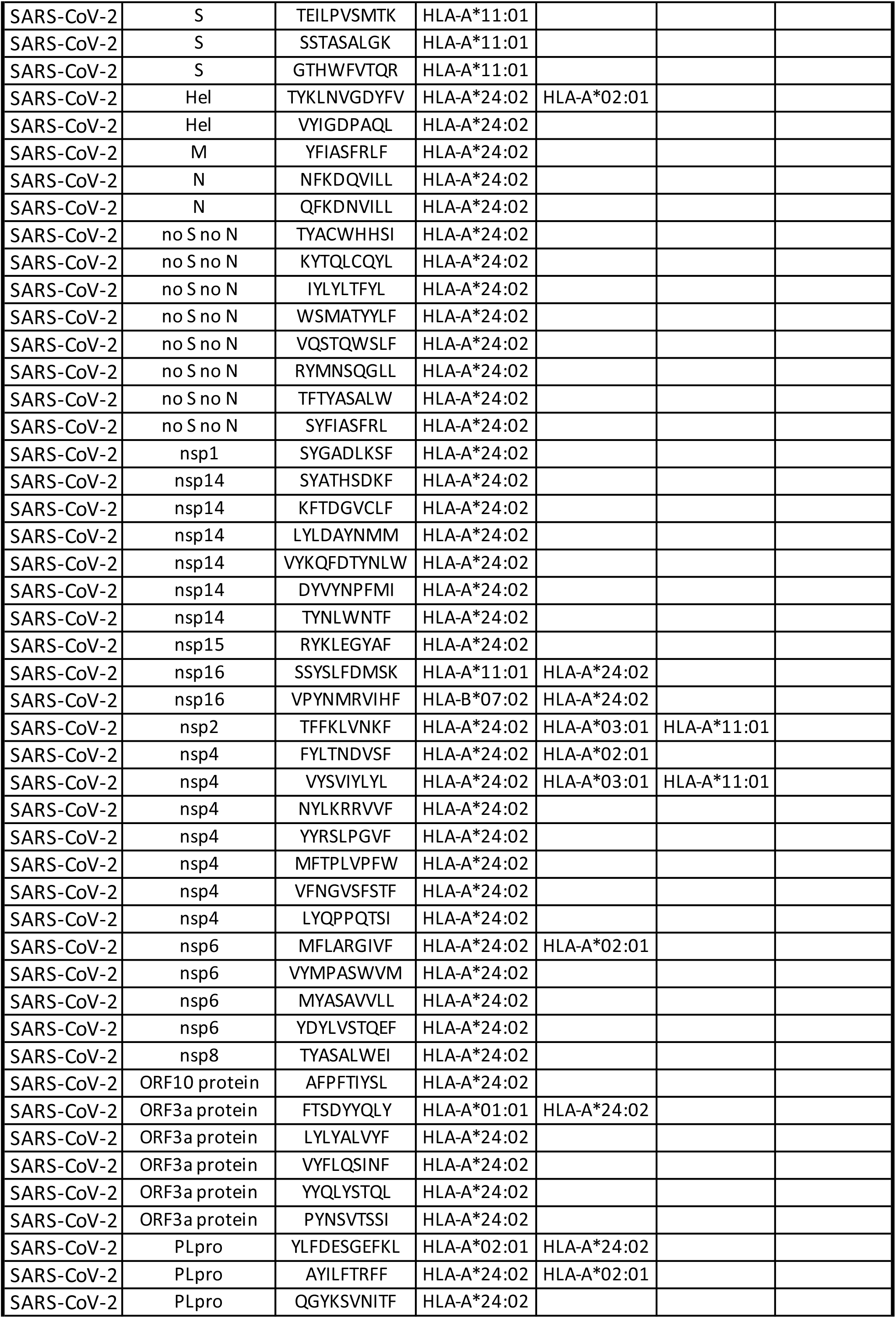
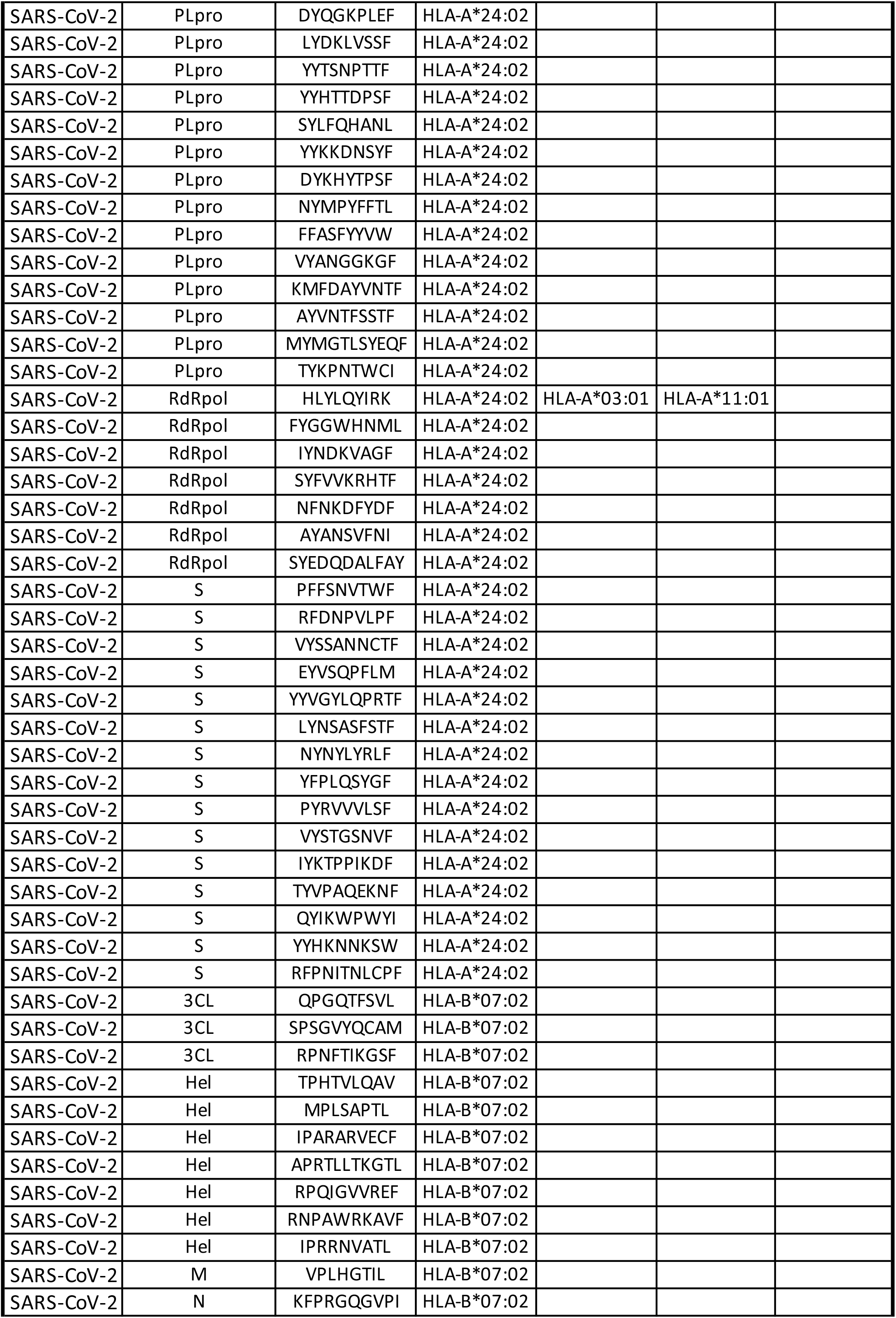
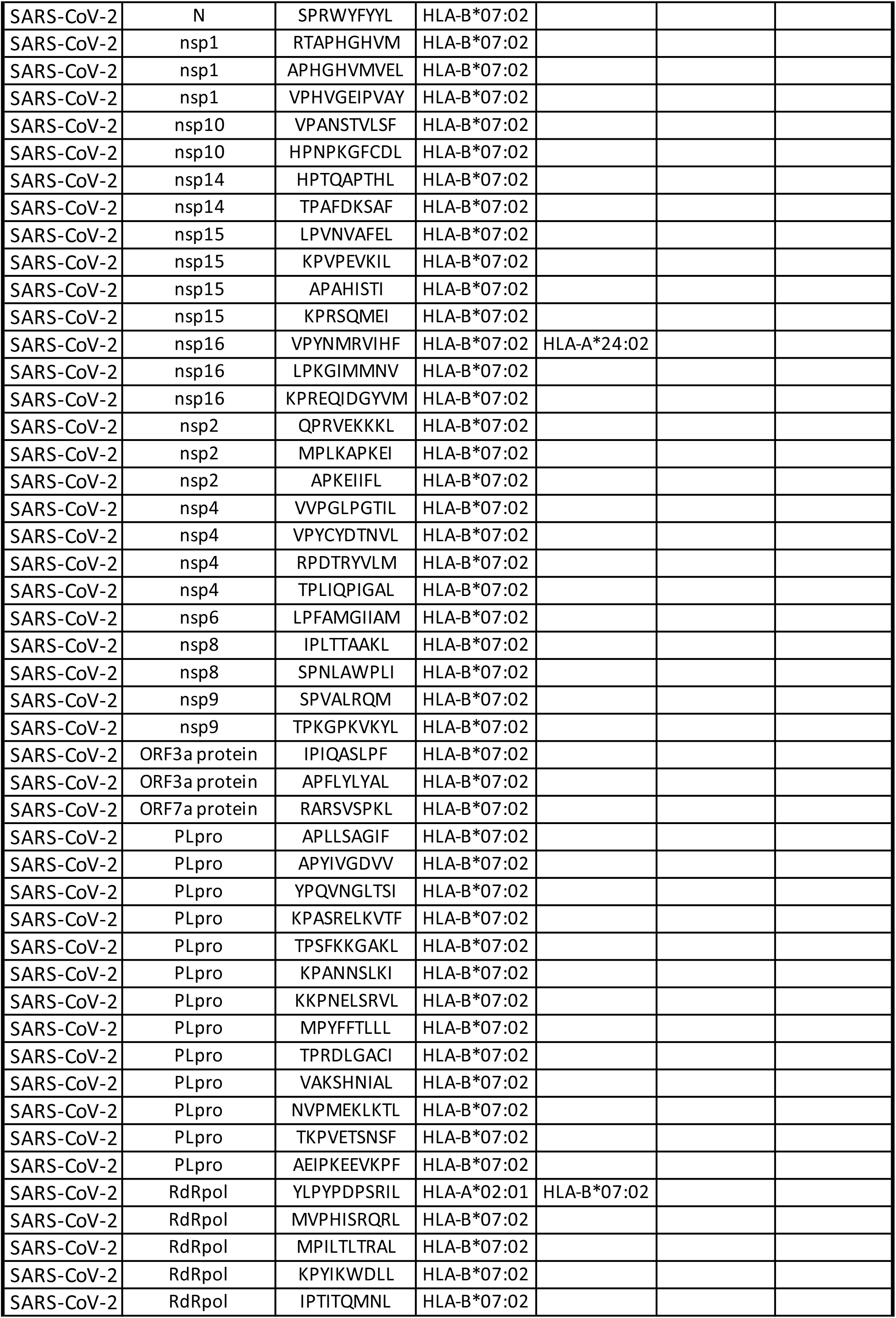
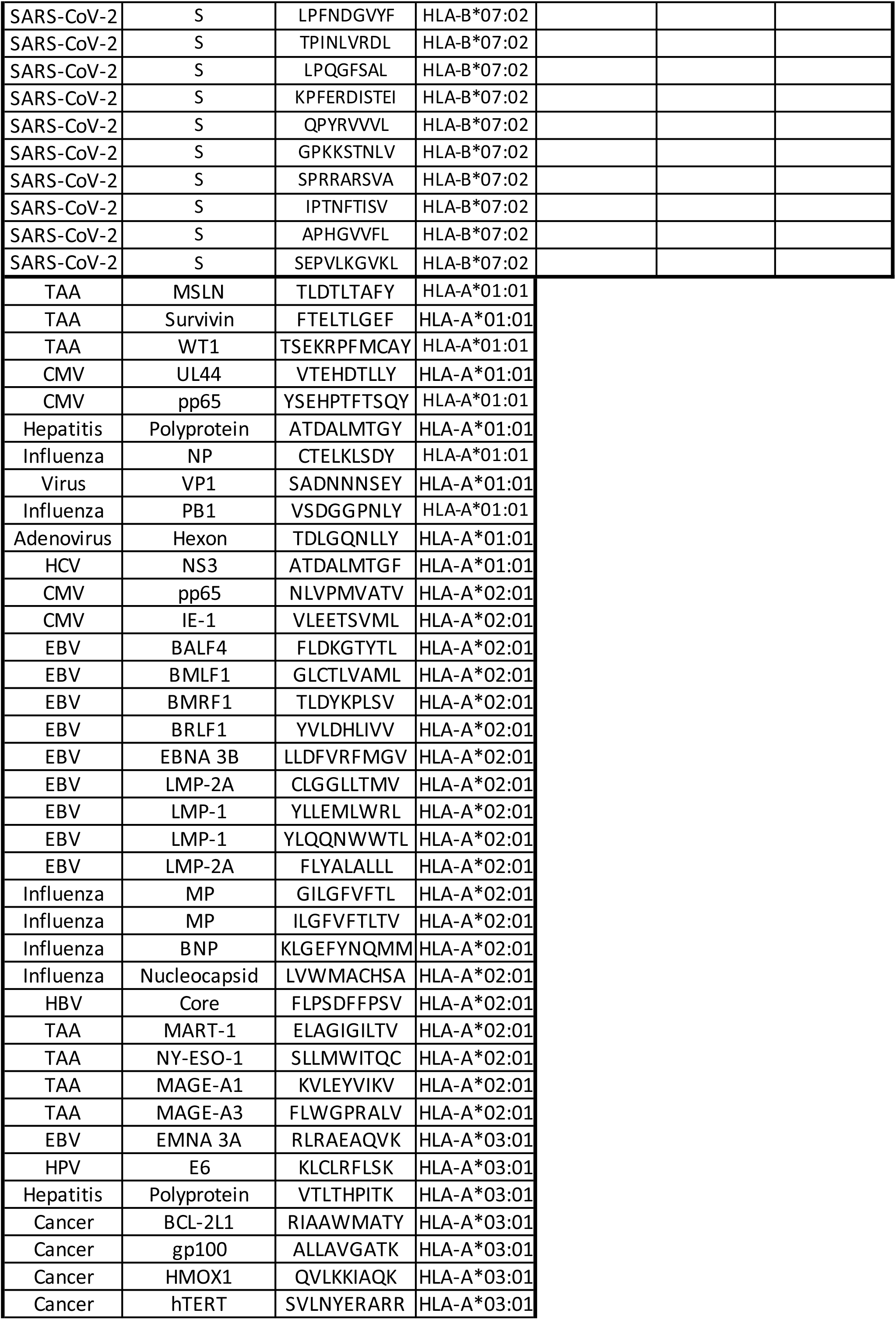
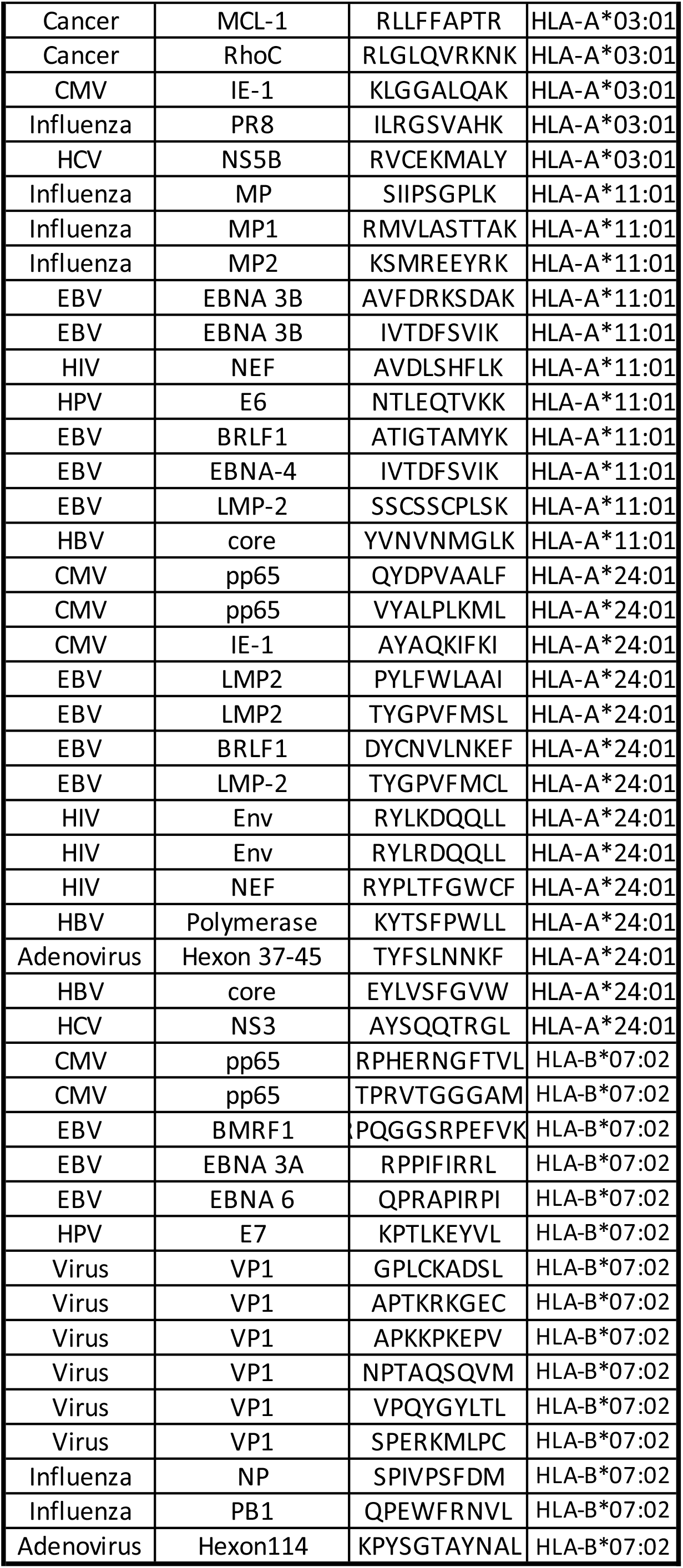

**Table S4.**
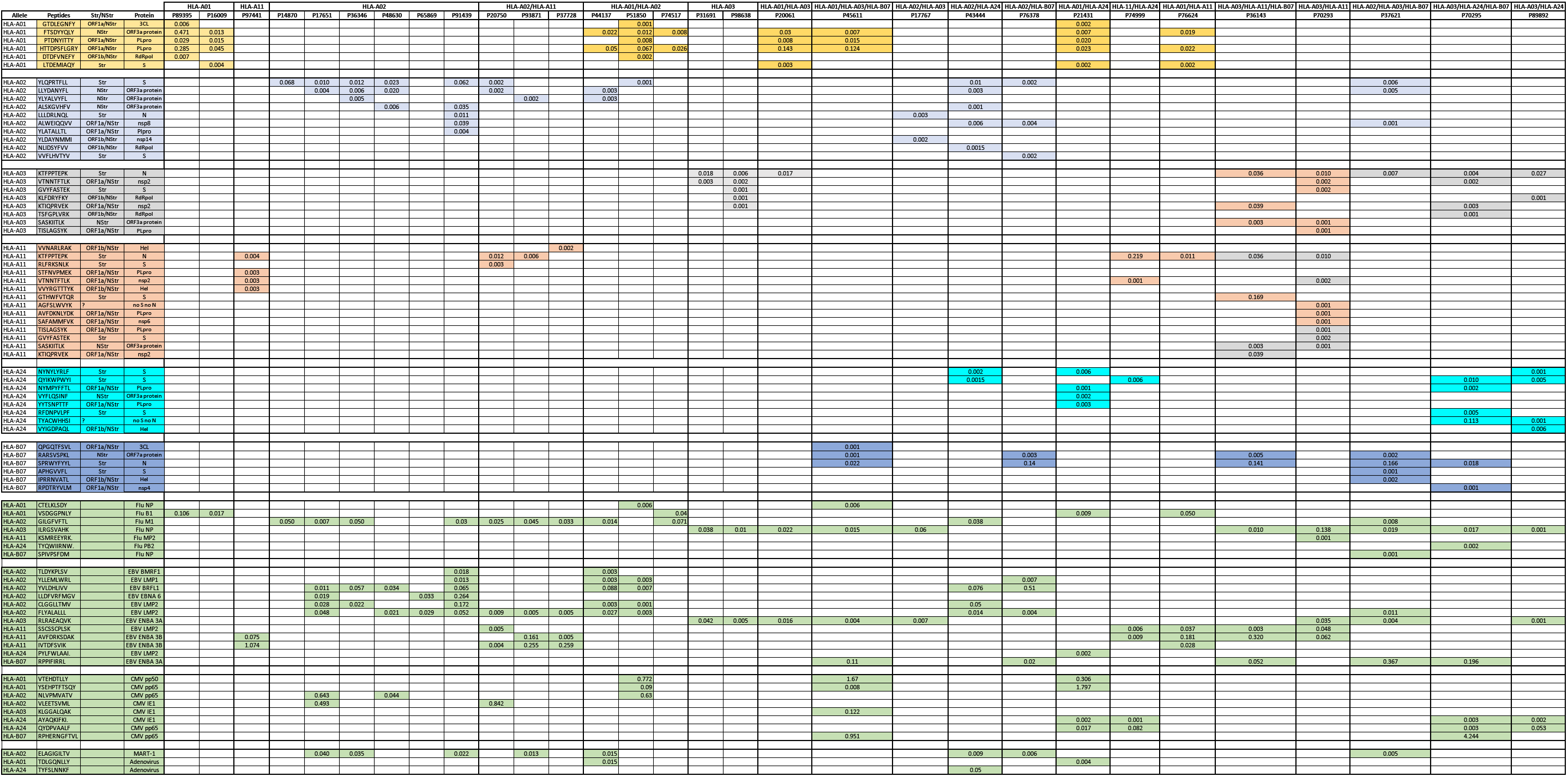

**Table S5.**

| **Metal** | **Antibody** | **Clone** | **Provider** |
| --- | --- | --- | --- |
| Y-89 | CD45 | HI30 | *Fluidigm* |
| Cd-112/114 | CD14 | TUK4 | *Invitrogen* |
| Cd-112/114 | CD19 | SJ25-C1 | *Life Technologies* |
| In-115 | CD57 | *HCD57* | *Biolegend* |
| Pr-141 | CD103 | *B-Ly7* | *eBiosciences* |
| Nd-142 | CD56 | *NCAM 16.2* | *BD* |
| Nd-143 | HLA-DR | *L243* | *Biolegend* |
| Nd-144 | CD3 | *UCHT1* | *Biolegend* |
| Nd-145 | CD4 | *SK3* | *Biolegend* |
| Nd-146 | CD8 | *SK1* | *Biolegend* |
| Sm-147 | CD45RA | HI100 | *Biolegend* |
| Nd-148 | CD45RO | *UCLH1* | *Biolegend* |
| Sm-149 | CD160 | MAB6700 | R&D |
| Nd-150 | Granzyme B | *CLB-GB11* | *ABCAM* |
| Sm-152 | CD38 | *HIT2* | *Biolegend* |
| Eu-153 | KLRG1 | *13F12F2* | *eBiosciences* |
| Gd-155* | CLA | *HECA-452* | *Biolegend* |
| Gd-158 | CD27 | *LG.7F9* | *eBiosciences* |
| Tb-159 | CXCR3 | MAB160-100 | R&D |
| Dy-161* | CD161 | *HP-3G10* | *Biolegend* |
| Dy-162* | CD95 | DX2 | *Biolegend* |
| Dy-163* | CD127 | *A019D5* | *Biolegend* |
| Er-168 | CCR7 | *150503* | R&D |
| Er-170 | CD244 | *C1.7* | *Biolegend* |
| Yb-173* | CD28 | *CD28.2* | *Biolegend* |
| Yb-174 | CD71 | *CY1G4* | *Biolegend* |
| Yb-176 | CD39 | *A1* | *Biolegend* |
| Bi-209 | CD16 | 3g8 | *Fluidigm* |

**Table S6.**
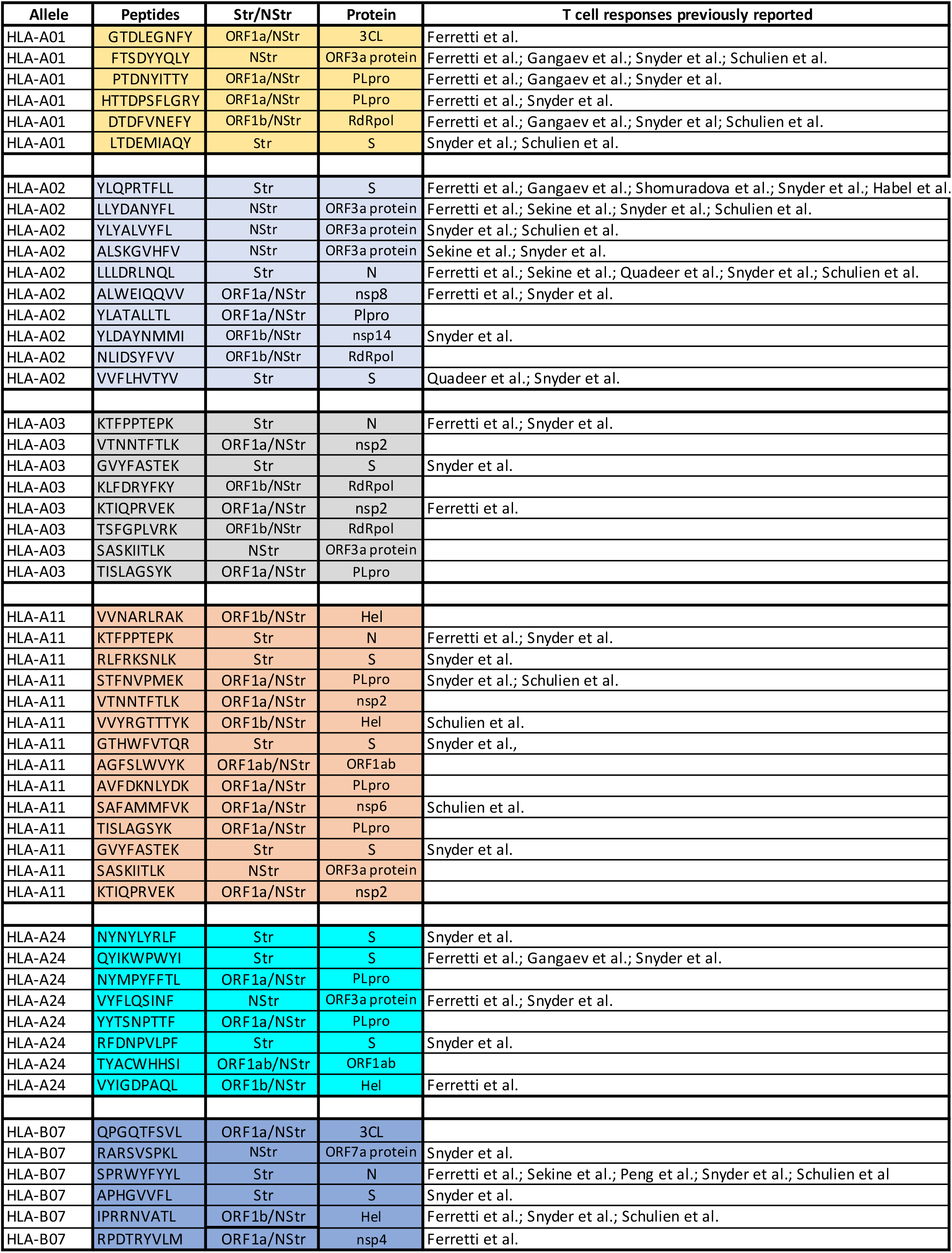

## References

1. Guan, W.-J. et al. Clinical Characteristics of Coronavirus Disease 2019 in China. N. Engl. J. Med. 382, 1708–1720 (2020).

2. Huang, C. et al. Clinical features of patients infected with 2019 novel coronavirus in Wuhan, China. The Lancet 395, 497–506 (2020).

3. Klein, S. et al. Sex, age, and hospitalization drive antibody responses in a COVID-19 convalescent plasma donor population. medRxiv (2020) doi:10.1101/2020.06.26.20139063.

4. Long, Q.-X. et al. Clinical and immunological assessment of asymptomatic SARS-CoV-2 infections. Nat. Med. 26, 1200–1204 (2020).

5. Braun, J. et al. SARS-CoV-2-reactive T cells in healthy donors and patients with COVID-19. Nature (2020) doi:10.1038/s41586-020-2598-9.

6. Grifoni, A. et al. Targets of T Cell Responses to SARS-CoV-2 Coronavirus in Humans with COVID-19 Disease and Unexposed Individuals. Cell 181, 1489–1501.e15 (2020).

7. Le Bert, N. et al. SARS-CoV-2-specific T cell immunity in cases of COVID-19 and SARS, and uninfected controls. Nature 584, 457–462 (2020).

8. Peng, Y. et al. Broad and strong memory CD4 + and CD8 + T cells induced by SARS-CoV-2 in UK convalescent individuals following COVID-19. Nat. Immunol. 1–10 (2020) doi:10.1038/s41590-020-0782-6.

9. Sekine, T. et al. Robust T cell immunity in convalescent individuals with asymptomatic or mild COVID-19. Cell (2020) doi:10.1016/j.cell.2020.08.017.

10. Ni, L. et al. Detection of SARS-CoV-2-Specific Humoral and Cellular Immunity in COVID-19 Convalescent Individuals. Immunity 52, 971–977.e3 (2020).

11. Folegatti, P. M. et al. Safety and immunogenicity of the ChAdOx1 nCoV-19 vaccine against SARS-CoV-2: a preliminary report of a phase 1/2, single-blind, randomised controlled trial. The Lancet 396, 467–478 (2020).

12. Yu, J. et al. DNA vaccine protection against SARS-CoV-2 in rhesus macaques. Science 369, 806–811 (2020).

13. Zhu, F.-C. et al. Safety, tolerability, and immunogenicity of a recombinant adenovirus type-5 vectored COVID-19 vaccine: a dose-escalation, open-label, non-randomised, first-in-human trial. The Lancet 395, 1845–1854 (2020).

14. Grifoni, A. et al. A Sequence Homology and Bioinformatic Approach Can Predict Candidate Targets for Immune Responses to SARS-CoV-2. Cell Host Microbe (2020) doi:10.1016/j.chom.2020.03.002.

15. Prachar, M. et al. COVID-19 Vaccine Candidates: Prediction and Validation of 174 SARS-CoV-2 Epitopes. bioRxiv 2020.03.20.000794 (2020) doi:10.1101/2020.03.20.000794.

16. Fehlings, M. et al. Late-differentiated effector neoantigen-specific CD8+ T cells are enriched in peripheral blood of non-small cell lung carcinoma patients responding to atezolizumab treatment. J. Immunother. Cancer 7, 249 (2019).

17. Newell, E. W. et al. Combinatorial tetramer staining and mass cytometry analysis facilitate T-cell epitope mapping and characterization. Nat. Biotechnol. 31, 623–629 (2013).

18. Pittet, M. J. et al. High frequencies of naive Melan-A/MART-1-specific CD8(+) T cells in a large proportion of human histocompatibility leukocyte antigen (HLA)-A2 individuals. J. Exp. Med. 190, 705–715 (1999).

19. Mahnke, Y. D., Brodie, T. M., Sallusto, F., Roederer, M. & Lugli, E. The who’s who of T-cell differentiation: Human memory T-cell subsets. Eur. J. Immunol. 43, 2797–2809 (2013).

20. Elsaesser, H., Sauer, K. & Brooks, D. G. IL-21 Is Required to Control Chronic Viral Infection. Science 324, 1569–1572 (2009).

21. Kared, H., Fabre, T., Bédard, N., Bruneau, J. & Shoukry, N. H. Galectin-9 and IL-21 Mediate Cross-regulation between Th17 and Treg Cells during Acute Hepatitis C. PLOS Pathog. 9, e1003422 (2013).

22. Ferretti, A. P. et al. COVID-19 Patients Form Memory CD8+ T Cells that Recognize a Small Set of Shared Immunodominant Epitopes in SARS-CoV-2. medRxiv 2020.07.24.20161653 (2020) doi:10.1101/2020.07.24.20161653.

23. Gangaev A, K. P. Profound CD8 T cell responses towards the SARS-CoV-2 ORF1ab in COVID-19 patients. (2020) doi:10.21203/rs.3.rs-33197/v1.

24. Quadeer, A. A., Ahmed, S. F. & McKay, M. R. Epitopes targeted by T cells in convalescent COVID-19 patients. bioRxiv 2020.08.26.267724 (2020) doi:10.1101/2020.08.26.267724.

25. Schulien, I. et al. Ex vivo detection of SARS-CoV-2-specific CD8+ T cells: rapid induction, prolonged contraction, and formation of functional memory. bioRxiv 2020.08.13.249433 (2020) doi:10.1101/2020.08.13.249433.

26. Shomuradova, A. S. et al. SARS-CoV-2 epitopes are recognized by a public and diverse repertoire of human T-cell receptors. medRxiv 2020.05.20.20107813 (2020) doi:10.1101/2020.05.20.20107813.

27. Snyder, T. M. et al. Magnitude and Dynamics of the T-Cell Response to SARS-CoV-2 Infection at Both Individual and Population Levels. medRxiv 2020.07.31.20165647 (2020) doi:10.1101/2020.07.31.20165647.

28. Diao, B. et al. Reduction and Functional Exhaustion of T Cells in Patients With Coronavirus Disease 2019 (COVID-19). Front. Immunol. 11, (2020).

29. Neidleman, J. et al. SARS-CoV-2-specific T cells exhibit phenotypic features reflecting robust helper function, lack of terminal differentiation, and high proliferative potential. http://biorxiv.org/lookup/doi/10.1101/2020.06.08.138826 (2020) doi:10.1101/2020.06.08.138826.

30. Kared, H. et al. Immunological history governs human stem cell memory CD4 heterogeneity via the Wnt signaling pathway. Nat. Commun. 11, 821 (2020).

31. Pizzolla, A. et al. Resident memory CD8(+) T cells in the upper respiratory tract prevent pulmonary influenza virus infection. (2017) doi:10.1126/sciimmunol.aam6970.

32. Fehlings, M. et al. Checkpoint blockade immunotherapy reshapes the high-dimensional phenotypic heterogeneity of murine intratumoural neoantigen-specific CD8+ T cells. Nat. Commun. 9, 3000 (2018).

33. Finck, R. et al. Normalization of mass cytometry data with bead standards. Cytom. Part J. Int. Soc. Anal. Cytol. 83, 483–494 (2013).

34. Becht, E. et al. Dimensionality reduction for visualizing single-cell data using UMAP. Nat. Biotechnol. (2018) doi:10.1038/nbt.4314.

35. Levine, J. H. et al. Data-Driven Phenotypic Dissection of AML Reveals Progenitor-like Cells that Correlate with Prognosis. Cell 162, 184–197 (2015).

